# Fusiform Cells in the Dorsal Cochlear Nucleus Change Intrinsic Electrophysiological Properties and Morphologically Remodel Their Basal Dendrites with Age

**DOI:** 10.1101/2025.07.16.665173

**Authors:** Reginald J. Edwards, Michael R. Kasten, Kendall A. Hutson, Malcolm P. Lutz, Paul B. Manis

## Abstract

Age-related hearing loss (ARHL) is the most common cause of sensorineural hearing loss. The cochlear nucleus, the first central auditory structure to receive input from the cochlea, has been shown to be disrupted by ARHL. Fusiform cells (FC), the principal output cell of the dorsal part of the cochlear nucleus (DCN), mature physiologically during hearing onset. Specifically, FCs increase in rate of action potential (AP) rise and decay, stabilizing by postnatal day 14 (P14) in mice. However, whether FC intrinsic electrophysiological properties and morphological characteristics continue to change throughout the life of mice, and how they change due to ARHL, is unknown. We characterized electrophysiological and morphological properties of FCs from CBA/CaJ mice at five stages of age: preweaning (P15-20), pubescent (P21-49), young adult (P50-179), mature adult (P180-364), and old adult (P550-578). Our old adult mice had smaller auditory brainstem evoked response amplitudes and loss of some hair cells, indicative of ARHL onset. We observed no change in FC membrane properties with age. FCs from the old adult group had elevated firing rates, faster repolarization rates, and shorter AP half-widths. Morphologically, there was no change in FC soma shape or size. However, a significant decrease in basal dendritic arborization occurred between preweaning and pubescent ages, followed by an increase in our old adult group, suggesting age-dependent remodeling of the basal dendritic tree at the onset of ARHL. Together, these results suggest that FC physiology and morphology are relatively stable post weaning and become altered during the onset of ARHL.

**NEW & NOTEWORTHY:** *Ex vivo* patch-clamp recordings within the DCN are traditionally performed using young mice, rarely exceeding weaning age. Here, we were able to successfully record FCs from mice that were 15 days old up to 578 days old. We observed changes in FC firing properties, AP half-width, repolarization rate, and basal dendritic complexity in our old adult group, suggesting possible compensation for the development of age-related hearing loss.

## INTRODUCTION

Sensorineural hearing loss is the most common form of hearing impairment, and is characterized by pathology of the cochlea, auditory nerve, and central nervous system (1, 2). Age-related hearing loss (ARHL) is the most common cause of sensorineural hearing loss, affecting one-third of people aged 65-74 years and nearly half of those aged > 75 years in the United States alone (3). While considered by many to be a typical result of aging, ARHL has been linked to dementia, mood disorders, and overall poor quality of life (4), making it a critical research area. ARHL was first characterized through 4 etiologies each correlated with gradual hearing loss: loss of hair cells, atrophy of spiral ganglion cells and consequently the auditory nerve, atrophy of the stria vascularis, and structural changes in the basilar membrane (5–8). The CN is the first brain structure to receive and process acoustic information via auditory nerve fiber synapses from spiral ganglion cells of the cochlea. All auditory information traveling through the afferent auditory pathway must pass through the cochlear nucleus, making it an essential structure for understanding the effects of peripheral pathologies and central auditory system consequences due to ARHL. The cochlear nucleus is comprised of 2 main compartments: the ventral cochlear nucleus (VCN), and the dorsal cochlear nucleus (DCN). The DCN integrates auditory information with somatosensory (22–25), and vestibular (26, 27) signals that can contribute to accurate sound localization (28–30) and inhibition of self-generated sounds, even amid high background noise (31, 32). While there are multiple studies understanding the changes occurring in the VCN with ARHL (33–39), the effects of ARHL on the DCN are still being elucidated.

The functionality of the DCN is orchestrated by a network of inhibitory and excitatory neurons. Fusiform cells (FCs) are the major excitatory principal (projection) cell type in the DCN (40–46). FCs receive direct excitatory input from the auditory nerve fiber on their basal dendrites (47, 48) and excitatory input through granule cell parallel fiber stimulation to their apical dendrites (49–51). Excitatory descending inputs from the inferior colliculus and auditory cortex also reach the DCN (52–55). Additionally, FCs receive inhibitory inputs from other cells in the CN, including from cartwheel and tuberculoventral cells within the DCN and D-stellate cells of the VCN (56–63). These excitatory and inhibitory networks shape the FC responses to both narrow and wideband sounds. It has been shown via extracellular recordings that FCs from old anesthetized rats exhibit higher maximum noise-evoked and characteristic frequency-evoked firing rates compared to young anesthetized rats (14). While this finding highlights potential FC changes in response to ARHL, how the intrinsic electrophysiology or morphology of FCs is altered in response to ARHL is unknown.

Developmentally, the electrical excitability of FCs undergoes multiple changes from the onset of hearing (postnatal day 12-14 [P12-P14]) through at least P21 in murine models (64). Prominent changes include increased action potential amplitude, decreased half-width, faster rates of action potential rise, and increased depth of the afterhyperpolarization. While some neurons develop relatively mature firing properties by 2 weeks (65, 66), others are still immature at 2-3 weeks, continuing to mature their electrical signatures into adulthood (67–69). Whether mouse FCs continue to exhibit changes in electrophysiological and morphological characteristics after P21 is unknown.

In this study, we characterized the morphological and physiological properties of DCN FCs from shortly after hearing onset until the early stages of ARHL. We observed a decrease in auditory brainstem response amplitude with age, and a slight increase in threshold in old (up to P578) adult mice. While FC membrane properties were generally consistent from P21 through P578, we observed shorter AP half-widths, faster repolarization rates, and increased firing rates in our old adult FCs. Morphologically, we observed possible pruning of the basal dendrites between preweaning and pubescent FCs. Thereafter, basal dendritic branching gradually increases with age, returning to preweaning values in old adult FCs. This study shows that the properties and morphology of FCs are stable after early development and suggests that they may begin to show compensatory changes to loss of sensory activity at the onset of ARHL.

## MATERIALS AND METHODS

### Animals

CBA/CaJ mice of either sex from Jackson Labs (stock #000654), aged P15 to P578 were used in these experiments. Mice were group housed under a 12-hr light and 12-hr dark cycle in a facility room selected for low noise and minimal caretaker traffic. Hoods in the room were off when they were not in use. All experiments and procedures were done as approved by the University of North Carolina Internal Animal Care and Use Committee under protocols 21-123 and 24-083. The experimental design was to longitudinally examine the intrinsic excitability of DCN fusiform cells as a function of age. Mice were assigned to age groups, and are referred to as: preweaning (P15-20), pubescent (P21-49), young adult (P50-179), mature adult (P180-364), and old adult (P550-578).

### Auditory Brainstem Evoked Responses (ABR)

The hearing sensitivity of a subset of mice older than P20 was tested using click and tone ABR. For this test, mice were anesthetized with 100 mg/kg ketamine, 10 mg/kg xylazine intraperitoneally. When they ceased to respond to toe pinch, they were placed on a feedback heating pad with a rectal temperature probe to maintain body temperature at 37°C in a double-walled sound attenuating box made from 5/8” plywood, with inner and outer shells separated by 4”. The inner shell was lined with acoustic foam to reduce reflections. The eyes were covered with Optixcare eye lube to prevent dryness. Needle electrodes were placed at the vertex, right ear, and rump. ABR signals (and electrocardiogram [EKG]) were amplified by 10000X with a Grass P511J amplifier, filtered at 100-3000 Hz, and sampled with a Tucker Davis Technologies (Alachua, FL) RP2.1 digital signal processor at 97 kHz. Stimuli were digitally generated with a National Instruments NI6731 board at a 500 kHz sampling rate, utilizing a Tucker Davis PA5 attenuator to modulate intensity, then attenuated (6 dB) with a Tucker Davis SA5 speaker amplifier, to drive a TDT MF-1 free field speaker. The speaker was placed 5 cm directly in front of the tip of the mouse’s nose. The sound levels for each test frequency were calibrated against an ACO Pacific reference source (94 dB SPL, 995 Hz), using a ¼” ACO Pacific 7016 microphone (ACO Pacific, Belmont CA). During ABR recordings, the outer doors of the sound box were secured in place, and the door to the recording room was closed to minimize extraneous noise.

ABRs were acquired with a custom MATLAB program (ABR4) or in later experiments, with a custom Python program (pyabr3). In both cases, click stimuli were generated as 1.2 sec-long sweeps of 40 Hz, 100 µsec clicks, calibrated using the peak-to-peak method (70). Clicks were 100 µsec rectangular pulses. The peak-to-peak response (P1-N1) was measured using an automated peak detection algorithm (https://github.com/EPL-Engineering/abr-peak-analysis; peakdetect.py). This algorithm was incorporated into the analysis program to obtain the timing of responses and the latencies for the highest sound levels where there was a clear response. Responses were linearly extrapolated to lower (subthreshold) sound levels to obtain subthreshold noise values. The P1-N1 growth curve was fit to a Hill function, and the point where the curve exceeded 3 times the standard deviation of the baseline noise (averaged across traces) was taken as the threshold. Fits were visually verified, and the minimum latencies for peak detection were individually adjusted on a per-subject basis to ensure that the correct waveform components were identified.

### Slice Preparation

Mice were anesthetized with 100 mg/kg ketamine and 10 mg/kg xylazine (or given a supplemental dose if ABRs had just been measured), and transcardially perfused with an N-methyl-D-glutamine (NMDG) cutting solution (in mM: 93 NMDG, 30 NaHCO_3_, 25 glucose, 20 HEPES, 5 N-acetyl-L-cysteine, 5 ascorbic acid, 3 sodium pyruvate, 2.5 KCl, 2 thiourea, 1.2 KH_2_PO_4_, 0.5 CaCl_2_, 10 MgSO_4_, pH adjusted to 7.4 with HCl when oxygenated with 95% O_2_-5% CO_2_). This method has been used to improve cell survival in older mice (71). The brainstem was removed, blocked, and glued onto an agar “chair” (4% agar and 0.9% NaCl), and 250 µm slices were cut containing the DCN in the transstrial orientation (72) using a Campden 7000 smz-2 vibratome with ceramic blades (7550-1-C, Campden Instruments, Loughborough, England) in the same solution used for perfusion. Slices were incubated for 10 min at 35°C in the same NMDG cutting solution and then transferred to a HEPES-buffered saline (in mM: 92 NaCl, 30 NaHCO_3_, 25 glucose, 20 HEPES, 5 N-acetyl-L-cysteine, 5 ascorbic acid, 3 sodium pyruvate, 2.5 KCl, 2 thiourea, 1.2 KH2PO_4_, 2 CaCl_2_, 2 MgSO_4_, pH adjusted to 7.4 with NaOH when oxygenated with 95% O_2_-5% CO_2_) at room temperature for at least 1 hr before recording.

### Electrophysiological Recordings

Slices were placed into a chamber heated to 35°C (range of 34-36°C with a PH-2 heater, RG-26LP changer, Warner Instruments, Holliston, MA) and superfused with oxygenated artificial cerebrospinal fluid (ACSF) warmed by an in-line heater (TC-344B, Warner Instruments) at a flow rate of ∼2-3 mL/min. The ACSF consisted of (in mM): 134 NaCl, 3 KCl, 1.25 KH_2_PO_4_, 2 CaCl_2_, 1.3 MgSO_4_, 25 NaHCO_3_, 10 glucose, 2 Na-pyruvate, 3 myo-inositol, and 0.4 ascorbic acid. Slices and cells were visualized using oblique illumination under a Zeiss Axioskop 2FS microscope (Zeiss Instruments, Oberkochen, Baden-Württemberg, Germany), and images were obtained with a Retiga Electro R1 camera (Teledyne Photometrics, Tucson, AZ) utilizing digital image division for optimal visualization of DCN neurons. Recordings were obtained with borosilicate microelectrodes (BF150-86-10, Sutter Instruments, Novato, CA). Pipette resistances were 3-4 MΩ, and electrodes were not further treated to reduce pipette capacitance. The internal solution contained (in mM): 130 K-gluconate, 10 HEPES, 4 NaCl, 0.2 EGTA, 10 Tris phosphocreatine, 2 MgATP, 0.3 TrisGTP, pH to 7.2 with KOH, and osmolarity adjusted to 290 with H_2_O. The electrode solution included a saturating concentration of the fixable dye Lucifer Yellow CH-dipotassium salt (∼0.8%, Sigma-Aldrich, St. Louis MO, L1777) to identify the cells under fluorescence imaging during recording, and for subsequent morphological analysis. Undissolved Lucifer Yellow was removed from the electrode solution by centrifugation prior to filling the recording pipette. FCs were identifiable by their large elliptical soma, a generally bipolar arrangement of dendrites, and widely visible dendritic processes. Identification was confirmed by their morphological reconstruction and electrophysiological properties (59, 72–75). Whole-cell patch-clamp recordings were performed with a Multiclamp 700B amplifier (Molecular Devices, Sunnyvale, CA). Signals were low-pass filtered at 20 kHz and digitized at 100 kHz. Lower sampling rates failed to capture the maximal rising slope of action potentials accurately and lower filtering frequencies decreased the measured rising slope. Electrode resistance was cancelled during current injections by balancing the bridge. Pipette capacitance (typically 7-8 pF) was compensated ∼50% of the value measured in voltage-clamp (76) to 3.5-4.5 pF. While this compensation setting may not maximize the recording fidelity, it prevented overcompensation and ensured stability during the recording process. Data was acquired using the open-source program acq4 (77) (www.acq4.org). Tiled images of the slices in bright field (at low magnification, 4X air objective), and tiled and volume-stack images of the labeled cells *in situ* were taken with 10X and 40X water immersion objectives, with illumination from a 470 nm LED and a standard Zeiss FITC dichroic filter set, to help document the type and location of individually recorded cells.

### Electrophysiological Measurements

Cell responses to current injections from –200 to +200 pA (0.5 s duration, in 20 pA steps), –1 to 1 nA (1 s duration, in 50 pA steps), or 0 to 4 nA (1 s duration, in 200 pA steps) were recorded. The cell input resistance was computed from the maximum slope of a second-order polynomial fitted to the steady-state (mean from 0.4-0.5 s after the start of the current pulse) current-voltage relationship for currents in the range –1 to 0 nA. Resting membrane potential was computed from the 0.1 - 0.15 s baseline prior to the start of the current steps. The membrane time constant was measured from responses to small hyperpolarizing current steps that produced 2–10 mV voltage deflections, fit with a single exponential, and averaged across all traces that met criteria. The shapes of action potentials were also measured from the first spikes elicited by current steps at or just above rheobase, and which were followed by an interval of at least 50 ms before any following spike. Spike shape measures included spike threshold (measured as the first point on the rising phase of the action potential that exceeded a slope of 20 V/s), peak height (measured from threshold to action potential peak), spike half-width (measured at half the height from action potential threshold to peak), and afterhyperpolarization depth (measured from spike threshold to the nadir of the afterhyperpolarization). Several measures of firing rates in response to positive current injections were also taken. The adaptation index was measured for current steps that produced a firing rate between 80 and 120 Hz, by calculating the average of (st_i_ - st_i+1_) / (st_i_ + st_i+1_) where st_i_ indicates the time of the i^th^ spike in the train, and st_i+1_ is the time of the next spike in the train. This is the calculation used in the package *ipfx* from the Allen Institute (https://github.com/AllenInstitute/ipfx). Adaptation indices greater than 0 indicate decreases in the spike rate over time during current injections. Firing-rate versus current levels were computed and fit to a Hill function. If the firing rate was non-monotonic, only the rising part of the curve was used for the fit. The derivative of the Hill function was computed to obtain the steepest slope (gain) of the FI curve and the current at which that point occurred. The standard analysis used positive current steps from 0 to 1 nA, in 50 pA steps. In many cells, a second FI curve, with steps from 0 to 4 nA, in 0.2 nA steps, was also collected, and subject to the same analysis.

All electrophysiological measurements were computed using the Python suite “ephys” (www.github.com/pbmanis/ephys) running under Python 3.13. A standardized set of parameters were applied to all cells. Exclusion criteria included: cells with a holding current of more than −100 pA, junction potential-corrected resting potentials positive to - 50 mV, access resistances > 20 MΩ, noisy traces, or traces lacking sufficient data for certain calculations (such as time constants), to ensure that the data used for comparisons was not compromised by occasional poor recordings or cell health. All measures were selected from the recording protocols in each cell that were obtained with the lowest series resistance settings, to reduce voltage errors during current injections. If multiple acceptable runs met criteria in a cell, measurements were averaged. All traces are plotted with a correction for the junction potential of –12 mV.

### Immunofluorescent Staining

After electrophysiological recordings, slices were transferred to a 9% glyoxal/8% acetic acid/1X phosphate buffered saline (pH 7.4) fixation solution (78). Slices were incubated in fix overnight, then transferred to 1X PBS (21-040-CM, Corning) for long term storage. Subsequently, the slices were transferred to 24-well culture plates and placed on a mechanical shaker (IKA-Schuttler MTS 2, Wilmington, NC). Slices were rinsed once with fresh 1X PBS before being replaced with a 0.22 µm filter sterilized permeabilization buffer solution (0.6% Trition-100/1% Tween-20/1X PBS). Slices were subjected to rotation on the shaker at 100 rpm for 30-40 minutes at room temperature. After permeabilization, the solution was aspirated and the slices washed 3 times with 1X PBS before the addition of blocking buffer (4% Bovine Serum Albumin/1% Tween-20/1X PBS, 0.22 µm sterile filtered), then were left to shake at 100 rpm for 1 hr at room temperature. After the blocking step, slices were rinsed 3 times in 1X PBS before being incubated with rabbit-anti Lucifer Yellow (A5760, Invitrogen) at a 1:200 dilution in blocking buffer overnight at 4°C on a shaker at 100 rpm. After the overnight incubation, slices are rinsed three times in 1X PBS before the addition of donkey anti-rabbit 647 (A31573, Invitrogen) secondary antibody in blocking solution at a 1:500 dilution. Slices were covered with aluminum foil to protect the fluorophores from light, and incubated at room temperature for 1 hr, shaking at 100 rpm. After the secondary antibody incubation, slices were rinsed 3 times in 1X PBS and prepared for imaging.

Slices were transferred from the 24-well culture plate into a vial containing OptiMuS clearing solution (79) for 2 hrs in the dark at room temperature prior to imaging. Once the tissue was translucent, slices were transferred to a clean 22×40 mm rectangular No. 1.5 cover glass (CLS2980224, Corning/Sigma-Aldrich). The cover glass was prepared prior to transfer with the addition of 2 stacked 18×18 mm SecureSeal Imaging Spacers (654002, Grace Bio-Labs), each 0.12 mm thick, to provide 240 µm of imaging depth. Residual liquid from the transfer was aspirated and discarded. Once the slice was flat within the imaging spacers on the cover glass, one drop of SlowFade Gold Antifade Mounting Media (S36937, ThermoFisher Scientific) was added to the slice. Another clean cover glass was layered on top of the slice, creating a “sandwich” for imaging. Slices could be recovered from between the coverslips by injecting 150 µl of 1X PBS between the cover glasses and then deconstructing the cover glass. After imaging, slices were stored in 1X PBS at 4°C for future reimaging.

### Confocal Imaging Aquisition

Prepared cover glass ensembles imaged at the UNC Neuroscience Microscopy Core (RRID:SCR_019060) using a Zeiss 780 Laser Scanning Microscope. Zen Black (v2.3) software was used to define image acquisition parameters. A 633 nm Helium-Neon laser at 2-5% power was used to excite the secondary fluorophore to visualize Lucifer-yellow filled cells. All images were taken using a 63x oil immersion objective. Tiled, Z-stack images with 0.4-0.7 µm thick steps and a 57 µm wide pinhole were captured, encapsulating the entire DCN within a particular slice. Tiled images were stitched together with 10% overlap. A maximum intensity profile (MIP) image was generated from the stacked tiles for reference during morphological measurements.

### 3D Reconstruction and Analysis Using IMARIS

3D reconstructions of Lucifer-yellow filled FCs were created in IMARIS (v10.1-10.2) using confocal image stacks. MIP images were converted into IMARIS compatible format prior to reconstruction. Once imported, neurons were traced using the Filament tool. Somas were identified by localizing the fluorescence of a 3D box of approximately 20-30 μm in diameter. A soma model was then generated using machine learning algorithms, fitting the shape and complexity of a true soma. Next, seed points for the dendrites were generated, with sizes between 0.3-3.0 μm in diameter. Non-cellular seed points were filtered using local contrast thresholding set at 500. Selected seed points were manually curated as acceptable (following the fluorescent signal of the dendrite) or rejected (not a part of the neuron). This initial dataset was used as training data for the machine learning algorithm in IMARIS to predict the path of processes from the fluorescence image. This training and prediction occurred repeatedly until the skeleton of the neuron was visible. The skeleton was then converted to filaments. The same training-machine learning-prediction paradigm was iterated until the neuron was fully reconstructed. Manual filament additions, subtractions, and adjustments were made as needed. The reconstruction was split into apical and basal regions based on the directionality and morphology of dendrites. Dendrites extending into the molecular layer were categorized as apical. Dendrites extending towards the deep layer were categorized as basal. Primary dendrites protruding from the side of the neuron were classified as apical or basal based on the terminal point of the dendrite (molecular layer or deep layer). The whole cell, apical, and basal regions were then subjected to quantitative analysis with the IMARIS analysis functions. These functions generated a table of morphological measurements, along with a Sholl analysis of the dendritic tree structure.

### Cochlea Removal and Light Sheet Fluorescent Microscopy

Temporal bones containing the cochlea were harvested from a P573 mouse after brain removal and immersion fixed in 4% phosphate buffered paraformaldehyde. After 3-4 days, temporal bones were transferred to 10% EDTA for an additional 3 days, then returned to fixative. Cochleae were prepared for light sheet fluorescent microscopy (LSFM) after careful dissection and trimming away excess temporal bone. Each cochlea was rinsed overnight in PBS, then immersed in permeabilization solution (40% DMSO + 10% Triton X-100 in PBS, 37° C) for 2 days, then transferred to blocker solution overnight (5% bovine serum albumin + 5% Triton X-100 + 1% DMSO in PBS, 37° C). Cochleae were then exposed to primary antibodies for Myosin VIIa (1:200, Proteus Biosciences 25-6790, RRID: AB_10015251) and TuJ1 (1:100, BioLegend 801202, RRID: AB_2313773) in blocker at 37° C for 5 days; then washed 3 x 20 mins in PBS and transferred to species-specific Alexa Fluor tagged secondary antibodies (1:250 in blocker, 37° C) for 5 days. Cochleae were then washed in PBS (3 x 20 mins), dehydrated in ethyl alcohols (30, 50, 70, 90 and 100%) for 1 hr each, and fresh 100% ethyl alcohol overnight. Specimens were subsequently cleared with Spalteholz solution (5:3, methyl salicylate:benzyl benzoate) overnight, then stored in dibenzyl ether prior to imaging. Cochleae were imaged on a LaVision Ultramicrosope II (LaVision BioTec GmbH, Bielefeld, Germany) fitted with a sCMOS detector (Andor Neo 5.5 sCMOS camera), and images were imported into IMARIS (v10.1-10.2) for 3D analysis. Following previously described methodologies, a filament was traced along the basilar membrane (80). Once traced, a distance-to-frequency map was applied to the filament to assist in evaluating the degree of hair cell loss in AHRL along the basilar membrane (81).

### Statistics

Statistical analyses of electrophysiological measurements were performed with scripts in R (4.3.3, The R Foundation for Statistical computing Platform, 2024), using a 1-way ANOVA with age as the primary factor. When the test revealed significance, a Tukey post-hoc test was performed, comparing all age groups. Multiple cells were recorded per subject, so the subject was the observational unit (e.g., all measures in a single subject were averaged before statistical analysis). Plotted data points show individual cell data. Box plots show median, 25-75% confidence intervals, and whiskers show 5-95% confidence intervals.

Statistical analysis of morphological data was performed in GraphPad Prism (v10.2). 2-way ANOVA was performed across age groups, with age and Sholl distance from soma being the primary and secondary factors. Distance from the soma for cells in all age groups was capped at 250 μm for the basal dendrites and 200 μm for the apical dendrites and binned into 50 μm increments. When the test revealed significance across the age factor, a Tukey multiple comparisons test was performed as a simple effects within rows model (comparing branch number across age at each 50 μm bin). Least square (LS) means and studentized range statistics (q) were calculated for each age group bin comparison. Plotted Sholl data shows individual cell Sholl analysis via spaghetti line graphs with the mean points and SD superimposed on top of individual cell data. Gamma functions were fitted using the non-linear curve fitter in Prism, for each cell for the apical and basal dendrites separately. The gamma equation used in GraphPad Prism was 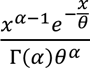. Initial values for alpha and theta were set to 2. Gamma curve fits generated measurements of skewness, variance, and smoothed peak branch number by distance measurements for each age group (Table 1,2). Skewness was calculated (in Excel) as 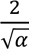 and the peak branch number by distance was calculated as *αθ*. Standard deviation of the peak branching number by distance was calculated in excel as 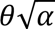. Gamma curves are plotted as solid lines, with shaded areas representing 95% confidence intervals. Cell surface area, volume, branch points, and terminal points were analyzed using a non-parametric Kruskal-Wallis 1-way ANOVA when data did not pass the Wilk-Shapiro normality test. If the data passed the Wilk-Shapiro test, a 1-way ANOVA with Brown-Forsythe correction was used. Data is plotted as box and whisker graphs with individual data points for each cell in an age group. Box plots show median, 25-75% confidence intervals, and whiskers show maximum and minimum values.

**Table 1:** Gamma curve fit analysis for the apical dendritic tree across ages.

|  | Preweaning | Pubescent | Young Adult | Mature Adult | Old Adult |
| --- | --- | --- | --- | --- | --- |
| Alpha | 5.32 | 5.23 | 5.23 | 5.16 | 5.25 |
| Theta | 32.83 | 22.13 | 23.83 | 22.28 | 24.25 |
| Degrees of Freedom | 3 | 3 | 3 | 3 | 3 |
| R squared | 0.79 | 0.94 | 0.94 | 0.98 | 0.97 |
| Peak Branching (Distance, $\mu\text{m}$) | 175 | 116 | 125 | 115 | 127 |
| Standard Deviation<br>(Distance, $\mu\text{m}$) | 76 | 51 | 54 | 51 | 56 |
| Skewness | 0.87 | 0.87 | 0.87 | 0.88 | 0.87 |

**Table 2:**
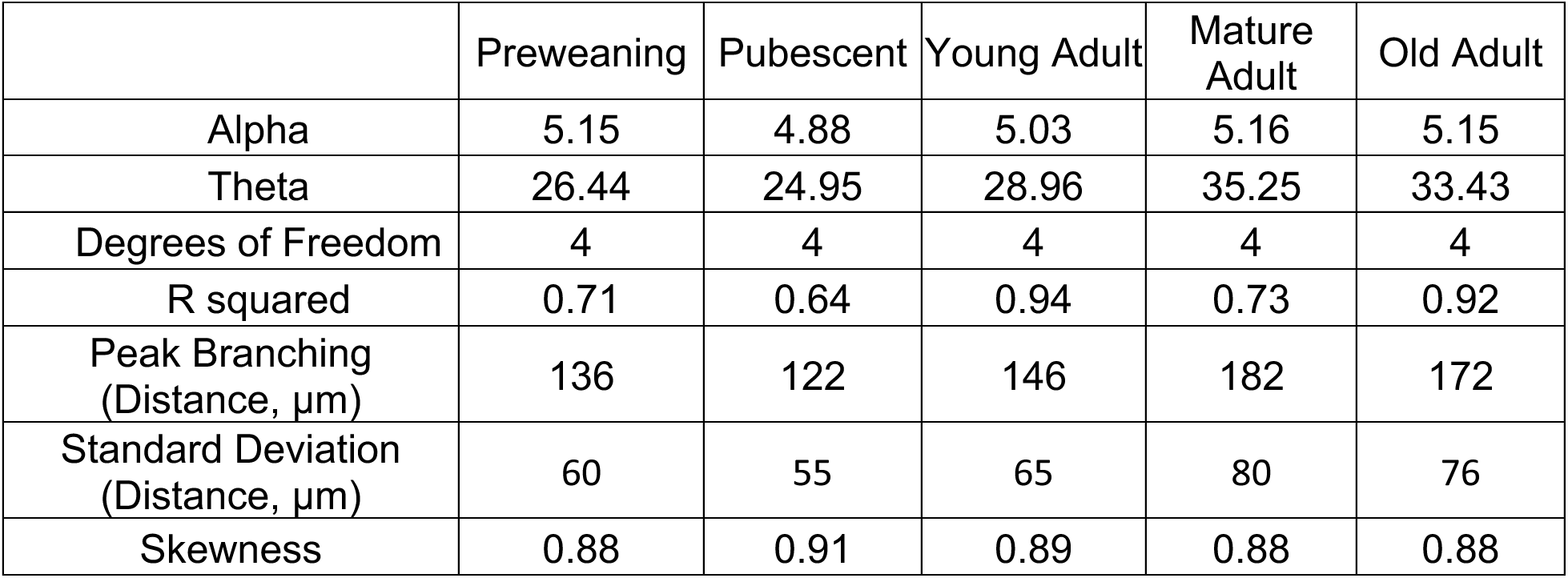
Gamma curve fit analysis for basal dendritic tree across age.

## RESULTS

### CBA/Caj Mice Demonstrate ARHL at 18 Months Through Auditory Brainstem Response Alterations and Hair Cell Loss

CBA/CAJ mice exhibit an age-dependent ABR threshold shift, specifically in the higher frequencies, making them useful candidates for studying age-related hearing loss (82). Threshold shifts begin around 18-20 months, with concurrent spiral ganglion cell degeneration and hair cell loss (82–84). This leads us to conclude that symptomatically, murine onset of ARHL appears to occur around 18-20 months of age in CBA/CAJ mice. To further refine this timeline, we recorded auditory brainstem responses to broadband clicks in mice across ages (except preweaning), 20-90 dB SPL (Fig.1A-C).

**Figure 1.**
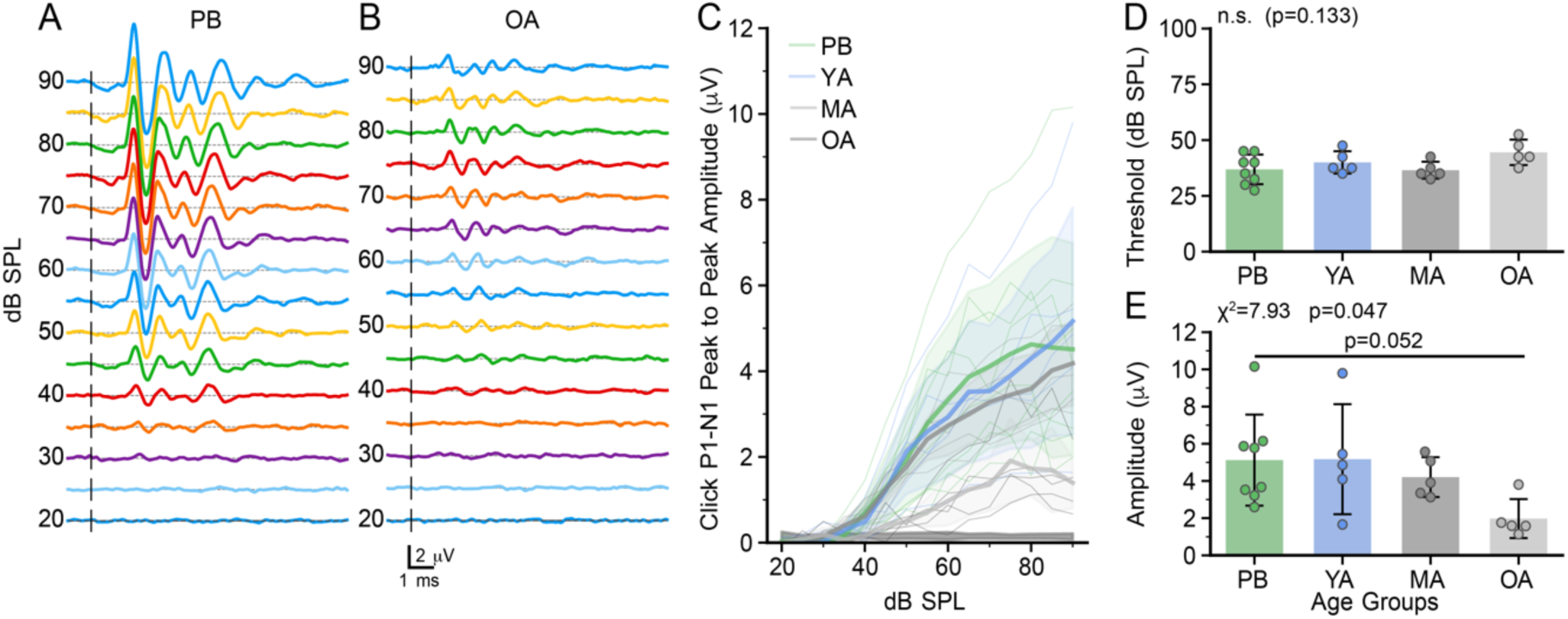
Click-evoked auditory brainstem evoked responses (ABR) as a function of age in a subset of the mice used in this study. *A-B:* ABRs as a function of level in a pubescent (*A*) and an old adult (*B)* mouse. *C:* Summary of individual ABR derived P1-N1 growth functions (thin lines) and means (thick lines). Shaded areas indicate the standard deviation of the data for each age group. *D:* Summary of ABR thresholds derived from the growth functions in C for each age group. *E:* Maximal amplitude of the P1-N1 amplitude for each age group. Bars indicate means, error bars indicate standard deviations. Categories: PB: pubescent, YA: young adult; MA: mature adult; OA: old adult.

There was a modest but not statistically significant (Kruskal-Wallis chi-squared = 5.60, df=3, p=0.133) threshold shift with broadband click stimuli with age, with a slight increase in the old adult group (Fig.1D). The auditory-nerve derived P1-N1 response amplitude showed a general age-related decline (Kruskal-Wallis chi-squared = 7.93, p=0.047; Dunn post-test with Bonferroni correction, old adult vs. pubescent p = 0.052; for all other comparisons, p > 0.1). The amplitude of the P1-N1 in the old adult group was 1.98 μV (SD 1.05), compared to mean values of 4.2 to 5.2 μV in the younger groups, suggestive of decreased auditory nerve input (Fig.1E).

To further assess changes occurring at the auditory periphery in old mice, hair cell distributions were observed using LSFM and mapped by their frequency representation along the basilar membrane in a old adult mouse (P573) (Fig.2A-C). Myosin VIIa is a vivid marker for hair cells (85). We observed hair cell loss all along the basilar membrane of our old adult cochlea, with the most extensive loss in the most apical and basal regions. Figure 2 E-G shows the constituent hair cells of the apical, middle, and basal 10% regions. In the apex (Fig2.D, 5.6 kHz to the apical terminus), IHC’s (Fig.2D, *hollow arrows*) appear disarrayed or missing. OHC’s (Fig.2D, *hollow arrowheads*) can be missing all rows (Fig.2D, *arrowhead* 1), reducing from 3 rows to 1 (Fig.2D, between *arrowheads* 2-3), with a poorly organized cluster just prior to the terminus (Fig.2D, *arrowhead* 4). In the middle turn, (Fig.2E, 16.5 – 22.0 kHz), hair cells appear relatively normal with only occasional missing IHC’s. Missing OHC’s are sometimes visible across all rows (Fig.2E, *arrowheads* 1 and 2) or as isolated OHC’s (Fig.2E, *arrowheads* 3 and 4). In the basal turn (Fig.2F, 58.3kHz to basilar membrane origin), a band of surviving IHC’s form a swath of cells near the origin, and a row of IHC’s re-appear just beyond the 58.3kHz mark. OHC are missing near the origin, then appear as a single cell (Fig.2F, *arrowhead* 1) before forming the characteristic three rows of cells (Fig.2F, *arrowhead* 2). IHC’s did not stain as intensely in the basal turn as adjacent OHC’s or IHC’s at other regions on the basilar membrane. Taken together with the functional ABR data, the old adult age range appears to demonstrate characteristic features of ARHL onset, with loss of auditory nerve input and loss of hair cells.

**Figure 2.**
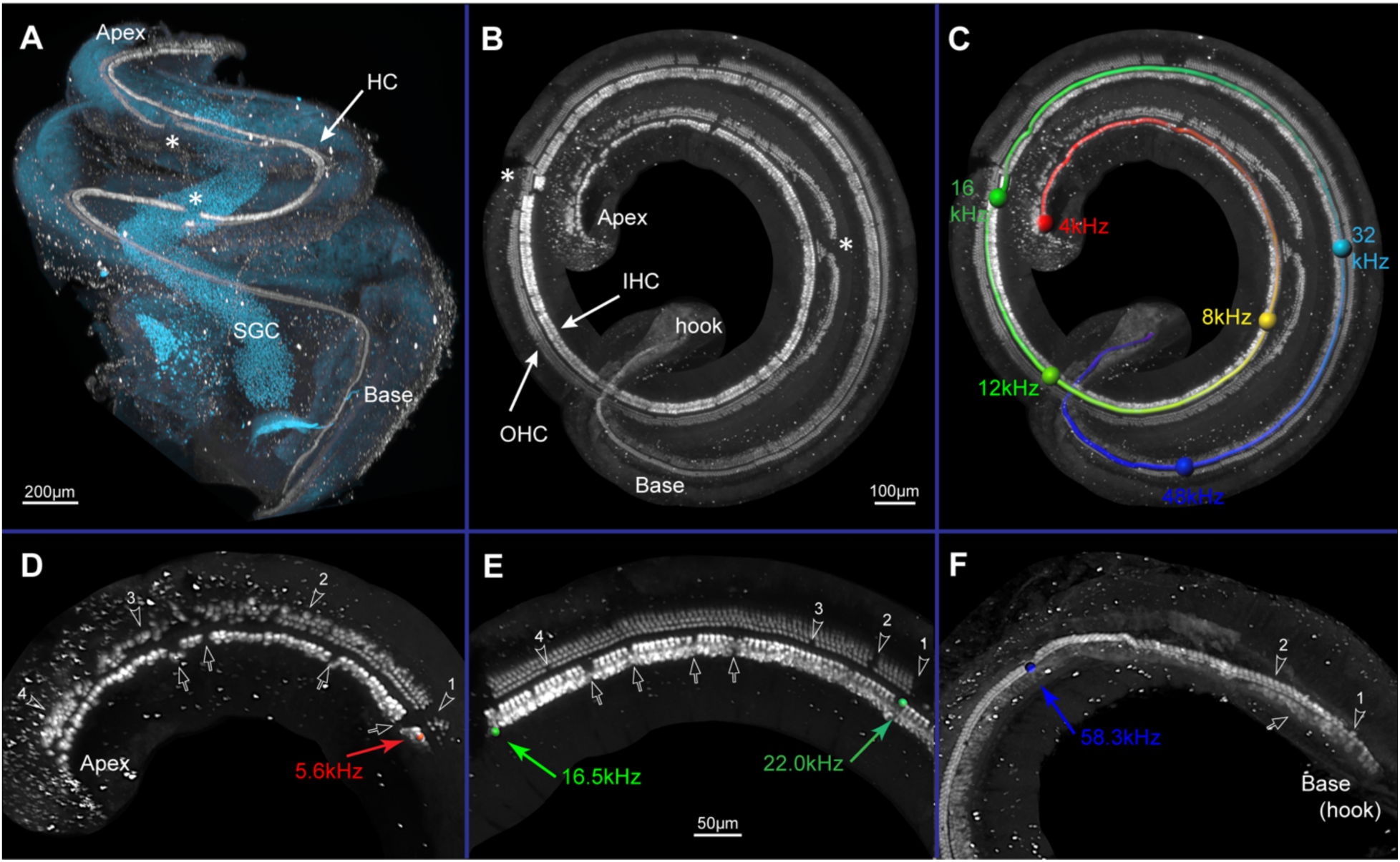
Light sheet fluorescence microscope images from a 573-day old mouse cochlea immunolabeled for Myosin VIIa. *A:* Three-dimensional view of labeled hair cells (HC) along basilar membrane from base to apex, spiral ganglion cells (SGC) imaged by autofluorescence. *B:* Virtually dissected basilar membrane (viewed from apex) showing labeled outer (OHC) and inner hair cells (IHC). *C:* Panel *B* overlayed with distance-to-frequency map. Spheres indicate pure tone frequency areas. *D-F:* hair cell distribution across apical (D), middle (E), and basal (F) 10% lengths of basilar membrane. Across all regions, IHC’s (*hollow arrows*) may appear disarrayed or missing; OHC’s (*hollow arrowheads*) can be missing all rows or reduced from three rows to one or two rows. *Asterisks* = artifact from errant forceps pinch during temporal bone dissection. *Scale bars*: for *C* same as in *B*; for *D-F* is in *E*.

### Passive Membrane Properties are Independent of Age

To assess changes in FC excitability as a function of age, whole-cell patch-clamp recordings were made from DCN FC in brain slices. There were no age-dependent changes in resting membrane potential (Fig.3A; 1-way ANOVA F(4, 158)=2.204, p=0.071, input resistance (Fig. 3B; 1-way ANOVA F(4, 152)=1.956, p=0.10) or membrane time constant (Fig.3C; 1-way ANOVA, F(4, 152)=0.428, p=0.79).

**Figure 3.**
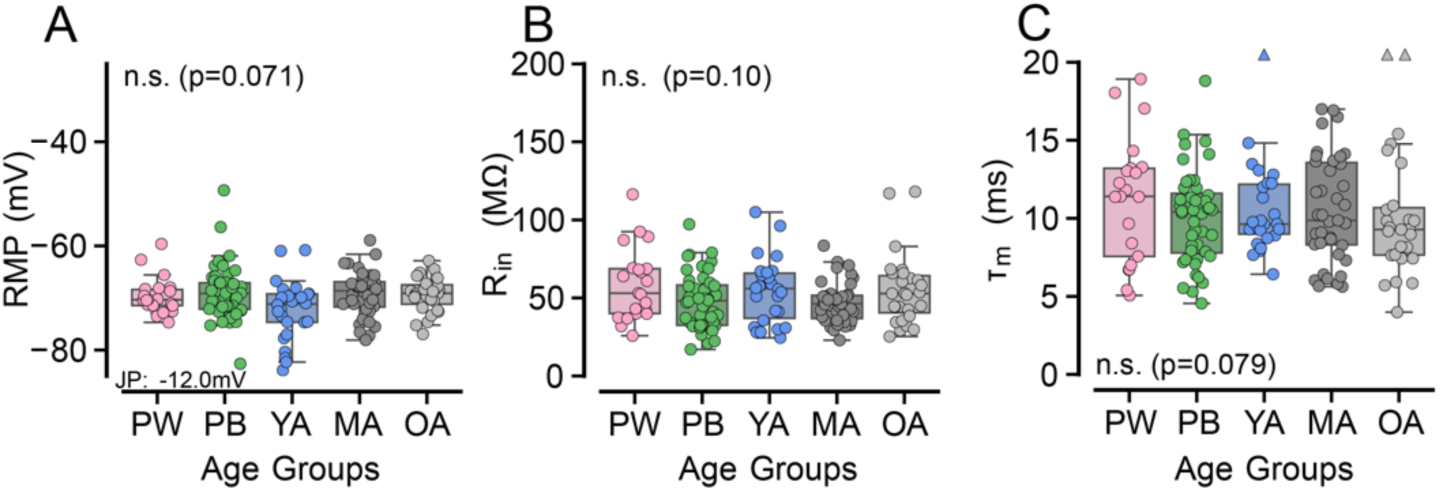
Passive membrane properties of fusiform cells across age. *A:* Resting membrane potential (RMP), corrected for the junction potential (JP) of −12 mV. Resting potential did not change with age. *B*: Cell input resistance (R_in_) was independent of age. *C:* The membrane time constant (τ_m_) did not change with age Age groups: PW: preweaning, PB: pubescent, YA: young adult, MA: mature adult, OA: old adult. Triangles at top of axes in *C* indicate measurements that are out of the range of the graph axes. Statistics indicate F and p for 1-way ANOVA; n.s. indicates not significant at the p=0.05 level.

### Fusiform Cells Repolarize Faster and Exhibit Narrower Action Potentials at the ARHL Onset Age

We next measured the shape of FC action potentials. Fig. 4A shows an example of first action potentials elicited at the lowest current step in a pubescent (green trace) and an old adult (light grey trace) neuron. The location of the action potential threshold and measurement of the afterhyperpolarization depth are indicated. Fig. 4B shows phase plots (dV/dt vs V) for these two action potentials, indicating the maximum rising and falling slopes, and the peak voltage. The red dots indicate the threshold and nadir of the AHP as shown in Fig. 4A. While there were no significant changes in maximum rate of action potential depolarization (Fig.4C: dV/dt rising: 1-way ANOVA F(4, 139)=1.654, p=0.16), the falling phase of the action potential (dV/dt falling, Fig. 4D) showed an age-related change (1-way ANOVA F(4, 139)=3.20, p=0.015), and post-hoc tests showed a significant difference between FCs from pubescent mice (441 (SD 106) mV/ms) and old adult mice (523 (SD 108) mV/ms; p = 0.017, Tukey method). Likewise, the ratio between the rising and falling slopes (dV/dt R/F ratio; Fig. 4E) showed an age-dependence (1-way ANOVA F(4, 139)=2.82, p=0.028), with post-hoc tests revealing a difference between the preweaning and mature adult FCs (preweaning: 1.61 (SD 0.30) vs mature adult; 1.43 (SD 0.19), p=0.024) as well as between the preweaning and old adult FCs (old adult: 1.43 (SD 0.22), p=0.036). Consistent with these measures, action potential half-width (Fig. 4F) showed a clear decrease with age (1-way ANOVA F(4, 139)=6.30, p=0.0011). Post-hoc tests revealed significance differences between the preweaning group (half-width 230 (SD 49) µs) and the mature adult group (half-width 201 (SD 28) µs; p=0.021), as well as between the preweaning and old adult age groups (half-width 186 (SD 30) µs; p=0.0002). The old adult FC action potentials were narrower than both FCs from pubescent mice (216 (SD 34) µs; p=0.0039) and from young adult mice (215 (SD 25) µs; p=0.015). These results indicate that with age, FC action potentials repolarize faster and consequently are narrower.

**Figure 4.**
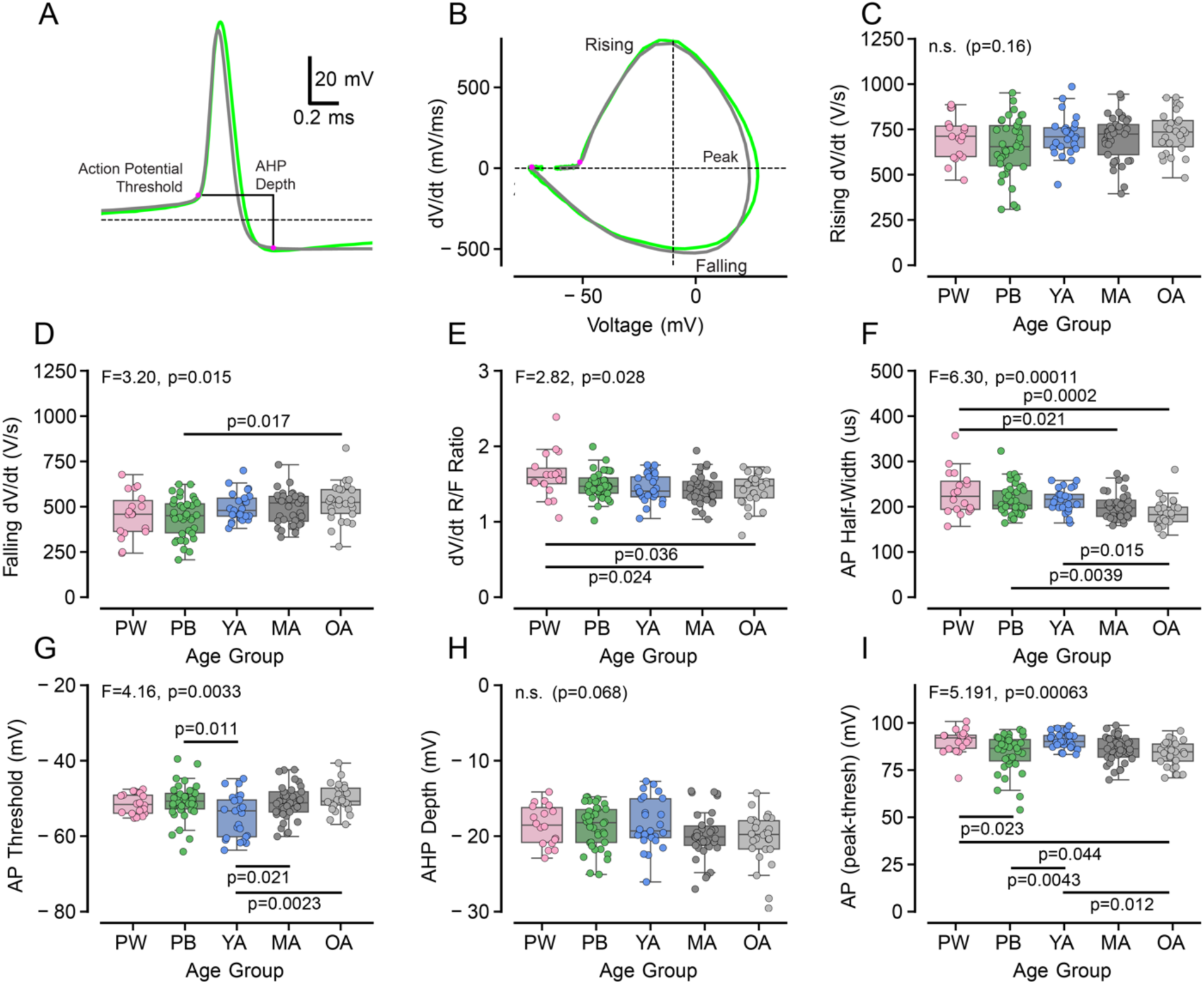
Individual action potential (AP) properties change with age. *A*: Representative APs from the pubescent (PB) and old adult (OA) age groups. The location of the AP threshold (20 mV/ms) and measurement of the afterhyperpolarization depth are indicated by magenta dots and the black line. Dashed line: −60 mV. *B*: Phase plots derived from the APs displayed in *A*. Labels show the maximal rising rate, falling rate, and location of the peak AP voltage. Magenta dots show the AP threshold and nadir of the AHP. *C-D:* Quantification of the maximum rising (*C*) and falling (*D*) slopes of APs across age. *E:* The ratio between the maximum rising and falling AP rates across age. *F-I:* Quantification of AP half-widths (F), threshold potential (G), afterhyperpolarization (H), and peak AP voltage (I) across age. Age groups: PW: preweaning, PB: pubescent, YA: young adult, MA: mature adult, OA: old adult. Statistics indicate F and p for 1-way ANOVA; n.s. indicates not significant at the p=0.05 level. Significant posthoc comparisons (Tukey corrected) are indicated by horizontal bars and p-values.

Action potential threshold (Fig.4G) showed age-dependence (1-way ANOVA F(4, 139)=4.16, p=0.0033), being slightly more negative in the young adult group (−54.7 (SD 5.7) mV) than in FCs from pubescent (−50.9 (SD 4.8) mV; p= 0.01), mature adult (−51.1 (SD 4.5) mV; p=0.021), and old adult (−50.0 (SD 3.7); p=0.0023) mice. Although the afterhyperpolarization appeared slightly deeper in the mature and old adult FCs (Fig. 4H), this was not significantly different (1-way ANOVA, F(4, 139)=2.243, p=0.068). Finally, the amplitude of the action potential, measured from threshold, was significantly different across ages (1-way ANOVA F(4, 139) = 5.191, p=0.00063). Post-hoc tests revealed differences between preweaning (90.3 (SD 6.7) mV) and pubescent age groups (83.8 (SD 9.8) mV; p=0.023), as well as between preweaning and old adult age groups (83.9 (SD 6.7) mV; p=0.044). Pubescent (83.8 (SD 9.8) mV) FCs had smaller action potentials than young adult FCs (90.7 (SD 4.4) mV; p=0.0043), while young adult FCs had larger action potentials than old adult FCs (p=0.012). However, the absolute peak amplitude of the action potential did not change across age (1-way ANOVA, F(4, 139)=2.242, p=0.068; see Supplemental Figure 1), suggesting that the peak height change was largely driven by changes in threshold. Taken together, these results indicate that the shape of FCs action potentials is altered at the onset of ARHL, with the predominant changes being a faster repolarization rate and a narrower AP half-width.

### Fusiform Cells Fire Faster at 18 Months

We next examined whether the intrinsic firing properties of FCs change with age. As expected from previous studies, FCs at all ages fired regular trains of action potentials with modest adaptation. Figure 5A shows the response of a fusiform cell from a P32 mouse to 1-sec long current pulses. At low current levels, firing can be brief and may cease before the end of the current pulse, but at higher levels, the firing was continuous throughout the trace. In response to hyperpolarizing current pulses, the membrane potential showed a sag typical of I_h_ currents and was usually followed by a short-lasting burst of spikes at stimulus offset. In some cases, there was a short delay to the first spike after hyperpolarization (blue trace in Fig.5A) as previously reported in guinea pigs and rats (74, 86), but this was rare in the cells from mice. Similar firing patterns were seen in our old adult age group (Fig.5B). The firing rate versus current curves for these 2 cells are shown in Fig.5C, along with the fits to a Hill function to the FI curve. The point at which the derivative of the FI curve (right axis) is maximal corresponds to the maximum firing gain. The arrows indicate the current levels at which the maximum gain is achieved. Fig.5D summarizes the FI curves across all FCs, coded by age. While the youngest 4 groups appear to cluster together, the old adult group has a slightly higher rate. To assess this, we examined the firing rate at 1 nA across ages (Fig.5E). A 1-way ANOVA revealed a significant effect of age (F(4, 148)=4.146, p=0.0033). Post-hoc tests showed that the rate of FCs from the old adult age group (340 (SD=80) Hz) was higher than in either the mature adult (288 (SD 74) Hz; p=0.036) or in the pubescent (270 (SD 60) Hz; p=0.0011) age groups.

**Figure 5.**
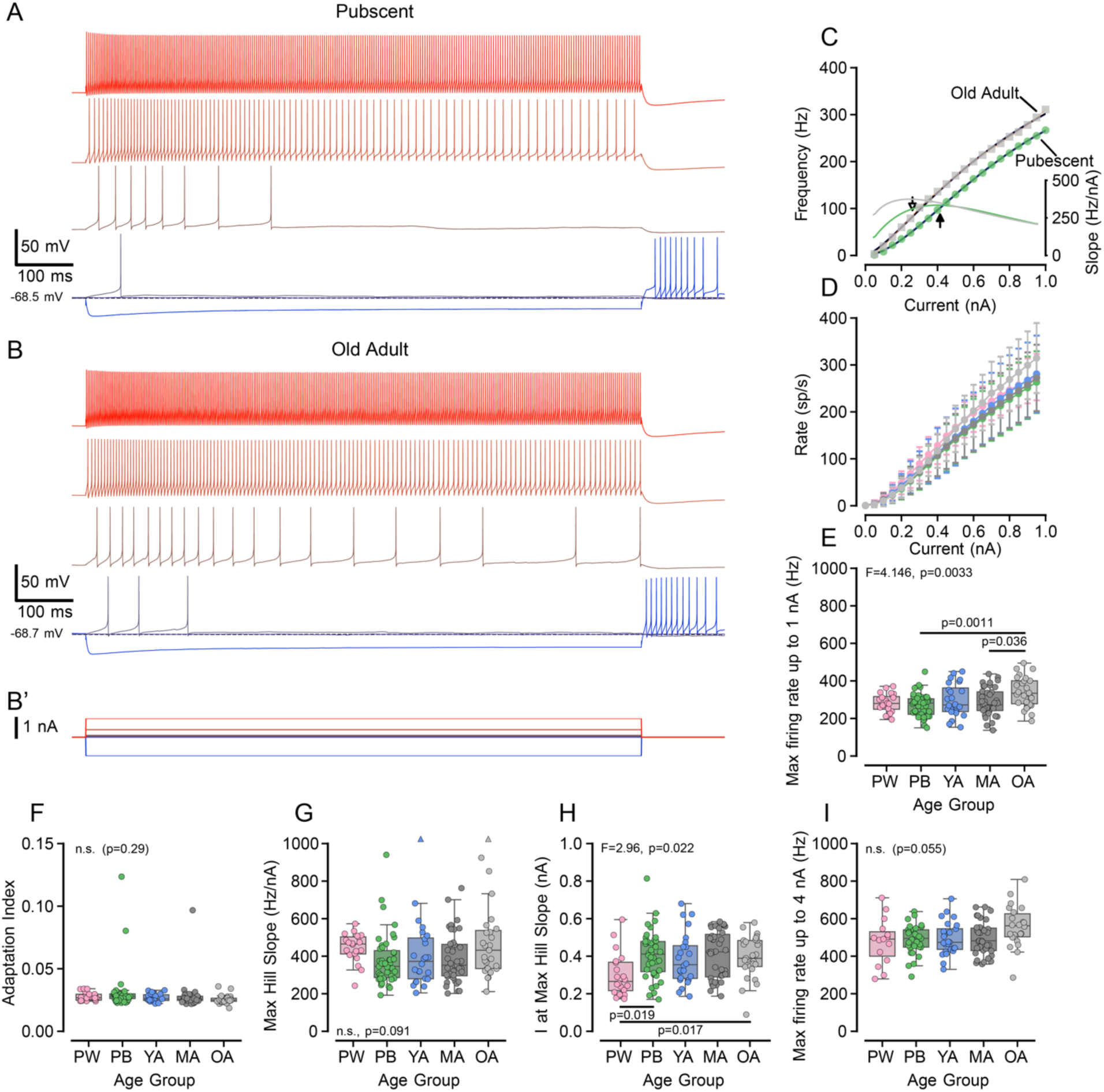
Fusiform cells change their firing properties with age. *A-B:* Example traces of responses to current injection from a pubescent (*A*) and an old adult (*B*) FC. Responses to a single hyperpolarizing current injection are also shown. *B’*: Current steps for *A* and *B*. *C:* Firing rate versus current injection for 1-s pulses from the cells in *A* and *B*. Left axis: firing rate is plotted with symbols with the smooth lines denoting best fit of a Hill function to the curve. Right axis: the slope (gain) of the Hill plot as a function of current. Arrows indicate the location of the maximal firing gain, as summarized in *G* and *H*. *D:* Comparison of firing rate curves for all groups, showing mean and 95% confidence intervals for each current level. *E-F*: Quantification of the maximum firing rate at 1 nA (*E*) and the rate of firing adaptation (*F*) across age groups. *G-I:* The maximal firing rate gain (*G*), current level at which the firing rate gain was largest (*H*) and the maximal firing rate at 4 nA (*I*) of FCs across age. Triangles at the top of the plot in panel G indicate measurements that are out of range of the graph axes. Age groups: PW: preweaning, PB: pubescent, YA: young adult, MA: mature adult, OA: old adult. Statistics indicate F and p for 1-way ANOVA; n.s. indicates not significant at the p=0.05 level. Significant posthoc comparisons (Tukey corrected) are indicated by horizontal bars and p-value.

The adaptation index, which measures how the firing rate changes over time, was not different between age groups (Fig.5F; F(4, 144)=1.25, p=0.29). We also evaluated the maximal firing rate gain by fitting the FI curves with a Hill function (as shown in Fig.5C) and extracting the maximum gain and the current at which the gain was maximum. The maximum gain of the FCs did not vary with age (Fig.5G, 1-way ANOVA, F(4, 148)=2.05, p=0.091). However, the current at which the gain was maximal did show an age-dependent increase (1-way ANOVA, F(4, 148)=2.96, p=0.22). The current level at which the gain was maximal for the preweaning age group (0.30 (SD 0.12) nA) was less than the pubescent (0.41 (SD 0.13) nA; p=0.019) and old adult (0.40 (SD 0.11) nA; p=0.017) age groups, suggesting slightly greater sensitivity to depolarizing currents in the preweaning age group. FC firing rates were also measured with a protocol that extended to 4 nA. The maximum firing rates at 4 nA did not quite reach statistical significance (1-way ANOVA, F(4, 123)= 2.382, p=0.055), likely due to the wide variation in firing rates at 4 nA in our preweaning age group. Remarkably, the mean firing rate for the old adult age group at 4 nA was 559 (SD 113) Hz of continuous firing, which is substantially higher than what is typically reported for FCs either *in vivo* or *in vitro*. It is likely that the narrower action potential half-widths and fast repolarization in the old adult FCs (Fig.4F) contribute to their ability to fire at such higher rates. Collectively, these results demonstrate an increase in the maximum firing rate of FCs from mice at the age of ARHL onset, and a decrease in sensitivity to injected currents after the end of the preweaning period (P21).

### Fusiform Cell Basal Dendrites Remodel Throughout Age

It was shown *in silico* that neuronal gain modulation can be influenced by dendritic morphology (87), and that neuronal firing properties can be influenced by dendrite geometry (88). To determine if the electrophysiological changes we observed with age were associated with morphological changes, recorded FCs filled with Lucifer-yellow were fixed, stained, imaged, and reconstructed in 3D (Fig.6). FCs have uniquely situated apical and basal dendrites (72, 89, 90); therefore, analysis was divided into apical and basal dendritic fields. The assignment of processes to the apical and basal dendritic fields was based on their orientation relative to the molecular layer (Fig.6, ML) or the deep layer (Fig.6, DL). Qualitatively, there was an abundance of dendrites, both apically and basally, in immature animals (Fig.6A). These dendritic arborizations appeared to simplify in the pubescent age group (Fig.6B), followed by an increase with age, most prominently in the basal dendrites (Fig.6C-E).

**Figure 6.**
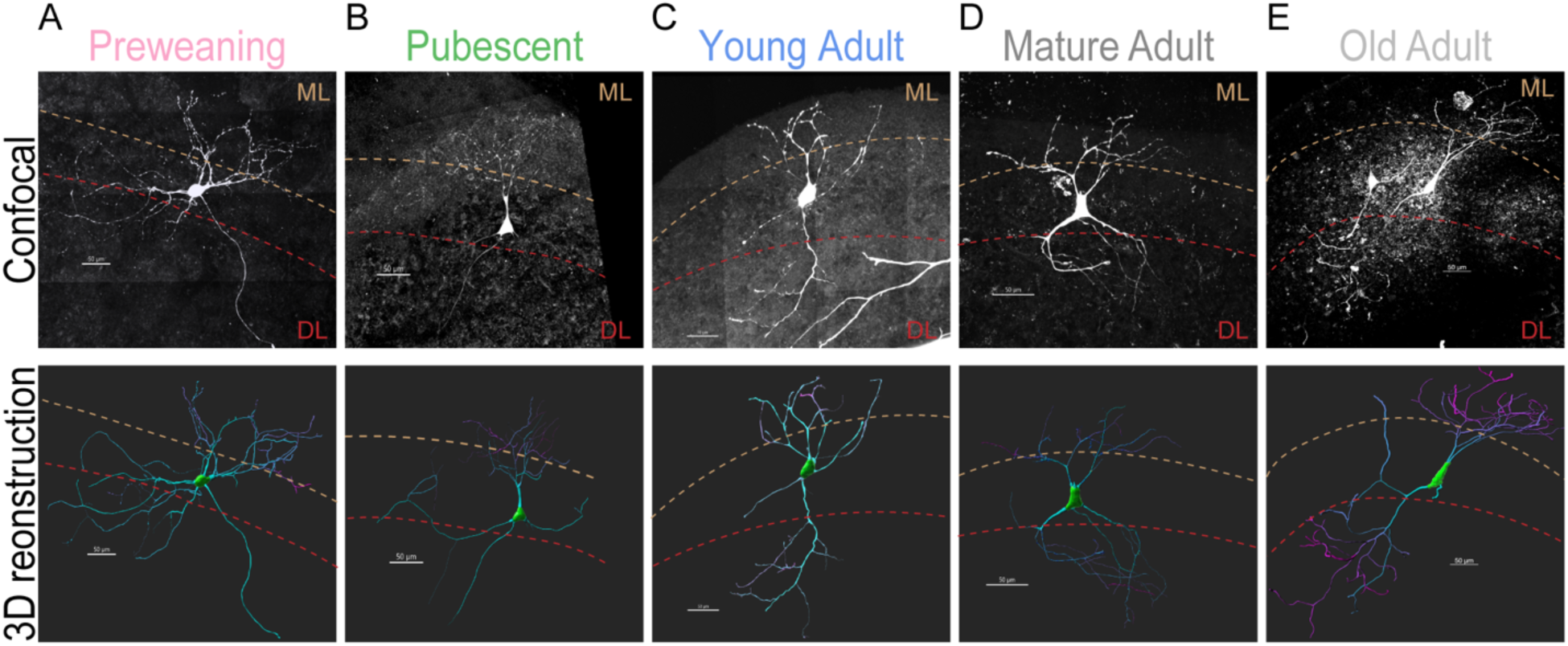
Representative fusiform cell confocal images and their corresponding 3D reconstructions. *A-E:* Confocal images (top row) are maximum intensity projections (MIPs) of lucifer-yellow injected FCs from brain slices, post electrophysiological recording. MIPs are then imported into IMARIS for 3D reconstruction (bottom row). Brown dashed line represents approximate bottom edge of the molecular layer (ML), while the red dashed line represents the approximate top edge of the deep layer (DL). Age groups: *A*: preweaning, *B*: pubescent, *C*: young adult, *D*: mature adult, *E*: old adult. All scale bars are 50 microns.

Subsequently, we measured the surface area and volume of FCs across age groups. We observed no change in the surface area (Fig.7A: Whole Cell field: Kruskal-Wallis 1-way ANOVA, p=0.72; Fig.7B: Apical dendritic field: Ordinary 1-way ANOVA with Brown-Forsythe, F (4, 42)=0.23, p=0.92; Fig.7C: Basal dendritic field: Kruskal-Wallis 1-way ANOVA, p=0.70), or cell volume (Fig.7D: Whole cell field: Kruskal-Wallis 1-way ANOVA, p=0.69; Fig.7E: Apical dendritic field: Kruskal-Wallis 1-way ANOVA, p=0.82; Fig.7F: Basal dendritic field: Kruskal-Wallis 1-way ANOVA, p=0.99) of FCs with age. These results are consistent with the lack of change in passive membrane properties.

**Figure 7:**
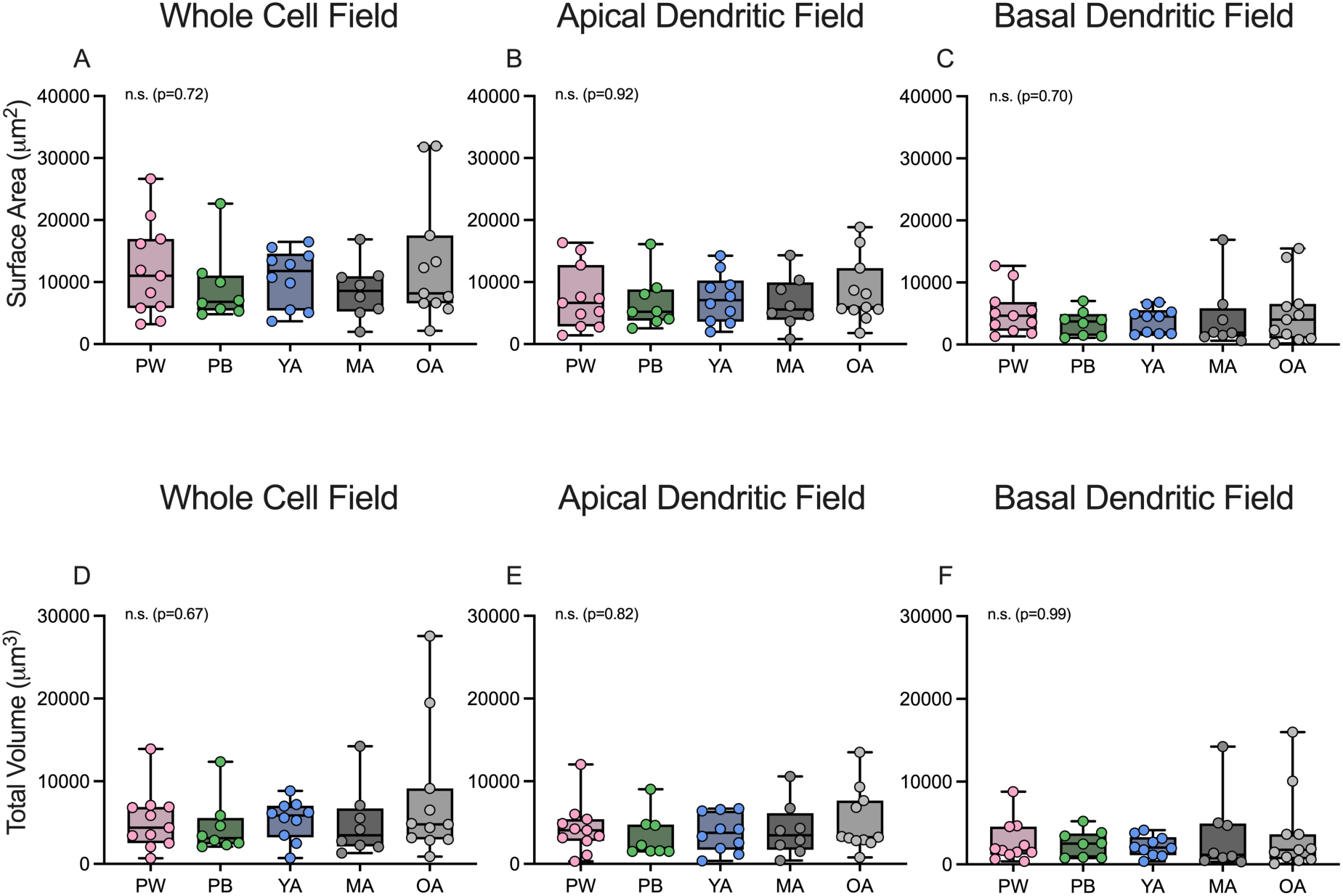
Fusiform cells do not change in size with age. *A-C*: Total surface area of the entire reconstructed neuron (*A*), the apical dendritic field (*B*), or the basal dendritic field (*C*). *D-F*: Total volume of the entire reconstructed neuron (*D*), the apical dendritic field (*E*), or the basal dendritic field (*F*). Either Kruskal-Wallis or Brown-Forsythe 1-way ANOVA was used for statistical analysis. n.s. = not significant (p values given in parentheses). Age groups: PW: preweaning, PB: pubescent, YA: young adult, MA: mature adult, OA: old adult.

We next set out to see whether overall terminal point or branch number were altered in FCs. There was no change in the overall number of terminal points (Fig.8A: Whole cell field: Kruskal-Wallis 1-way ANOVA, p=0.20; Fig.8B: Apical dendritic field: Kruskal-Wallis 1-way ANOVA, p=0.89; Fig.8C: Basal dendritic field: Kruskal-Wallis 1-way ANOVA, p=0.30) or branch points (Fig.8D: Whole cell field: Kruskal-Wallis 1-way ANOVA, p=0.21; Fig.8E: Apical dendritic field: Kruskal-Wallis 1-way ANOVA, p=0.82; Fig.7F: Basal dendritic field: Kruskal-Wallis 1-way ANOVA, p=0.28) of FCs with age. Interestingly, the basal dendrite branch and terminal point counts showed a small decrease from preweaning to pubescent age groups, followed by an increase as the FCs aged. This prompted us to look further into more robust ways of looking at dendritic complexity along the dendritic tree.

**Figure 8:**
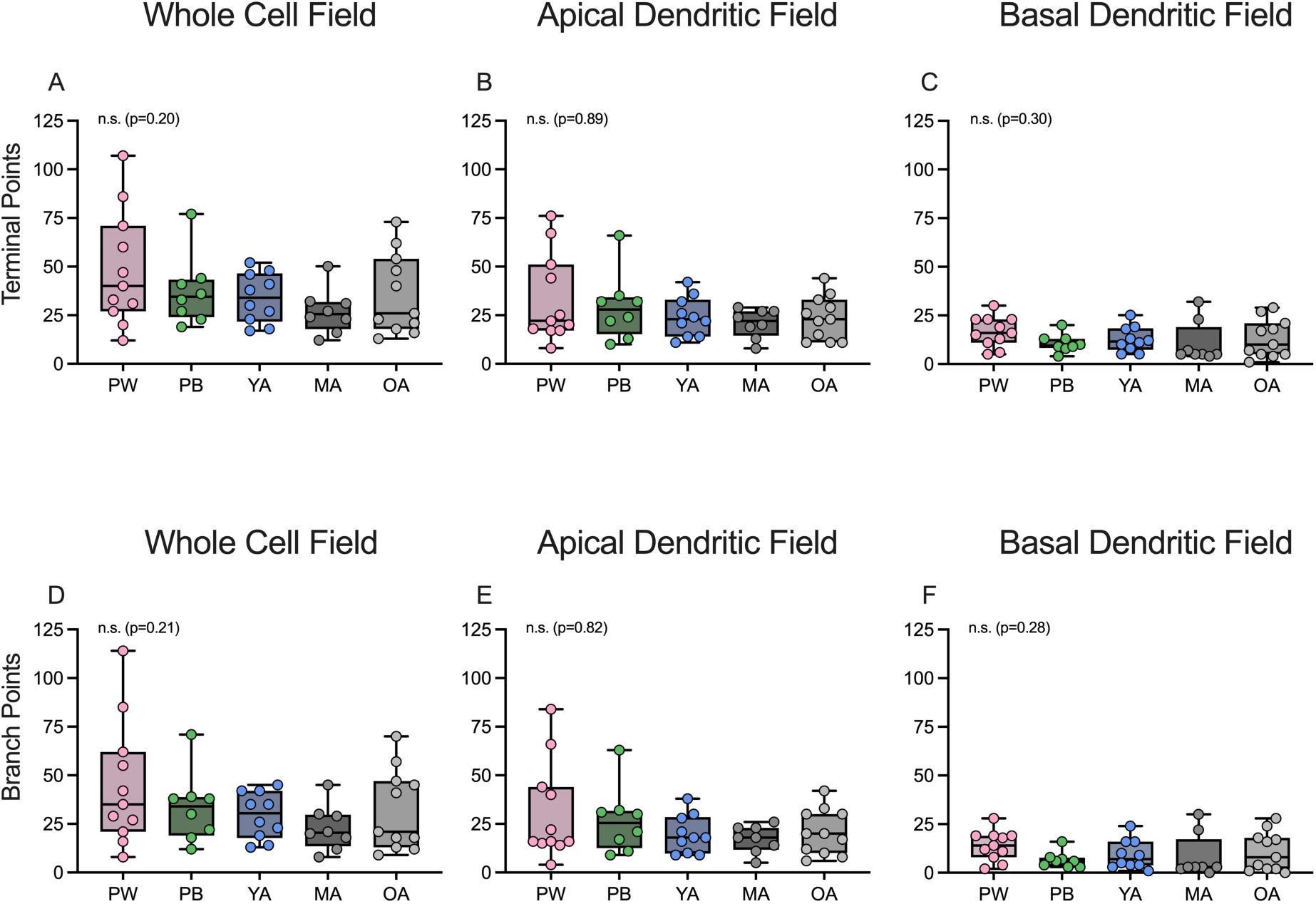
Fusiform cells do not change overall branching or terminal point number with age. *A-C*: Total terminal point number of the entire reconstructed neuron (*A*), the apical dendritic field (*B*), or the basal dendritic field (*C*). *D-F*: Total branch point number of the entire reconstructed neuron (*D*), the apical portion (*E*), or the basal portion (*F*). Kruskal-Wallis 1-way ANOVA was used for statistical analysis, with alpha = 0.05. n.s. = not significant (p values given in parentheses). Age groups: PW: preweaning, PB: pubescent, YA: young adult, MA: mature adult, OA: old adult.

A Sholl analysis was applied to both the apical and basal dendrites of FCs to better understand if dendritic complexity is altered with age. We observed no statistical change in the branching structure of the apical dendritic tree of FCs with age (Fig.9A; 2-way ANOVA, F(4, 182) =0.6881, p=0.60). However, at 100 µm from the soma, there was a significant decrease in dendritic arborization between the preweaning group and all other age groups (Fig.9C-D; 2-way ANOVA, F(4, 189) =3.69, p=0.006. Tukey’s multiple comparisons: Preweaning (LS=6.6) vs pubescent (LS=2.6), q=5.4, adjusted p=0.002; Preweaning (LS=6.6) vs young adult (LS=3.8), q=4, adjusted p=0.041; Preweaning (LS=6.6) vs mature adult (LS=3.3), q=4.1, adjusted p=0.034; Preweaning (LS=6.6) vs old adult (LS=3.8), q=4, adjusted p=0.041; d.f. for all comparisons=189). This reduction in basal dendritic arborization from the preweaning group is consistent with dendritic pruning following hearing onset. There was a trend towards increasing basal dendritic arborization at 150 µm from the soma in our old adult FCs compared to the pubescent FCs (Tukey’s multiple comparisons: old adult (LS=5.2) vs pubescent (LS=2.1), q=3.8, adjusted p=0.055, d.f.=189). This increase in basal dendritic remodeling is interesting, as it occurs further away from the cell body than where we see pruning (Fig.9D). Indeed, by fitting a gamma curve to the data, we observed changes in the peak branch intersection distances with age (Table 1 and 2). The FC apical dendrites exhibit peak branching further from the soma at the preweaning age group (175 (76 SD) µm) versus all other age groups (pubescent=116 (51 SD) µm; young adult=125 (54 SD) µm; mature adult=115 (51 SD) µm; old adult=127 (56 SD) µm). In contrast, the basal dendrites showed an age-related increase in peak branching distance from the soma, with the mature adult (182 (80 SD) µm) and old adult (172 (76 SD) µm) age groups having the furthest branch intersection peaks compared to the younger age groups (preweaning=136 (60 SD) µm; pubescent=122 (55 SD) µm; young adult=146 (65 SD) µm). Taken together, these results suggest early developmental pruning followed by enhanced basal dendritic arborization in FCs with age.

**Figure 9:**
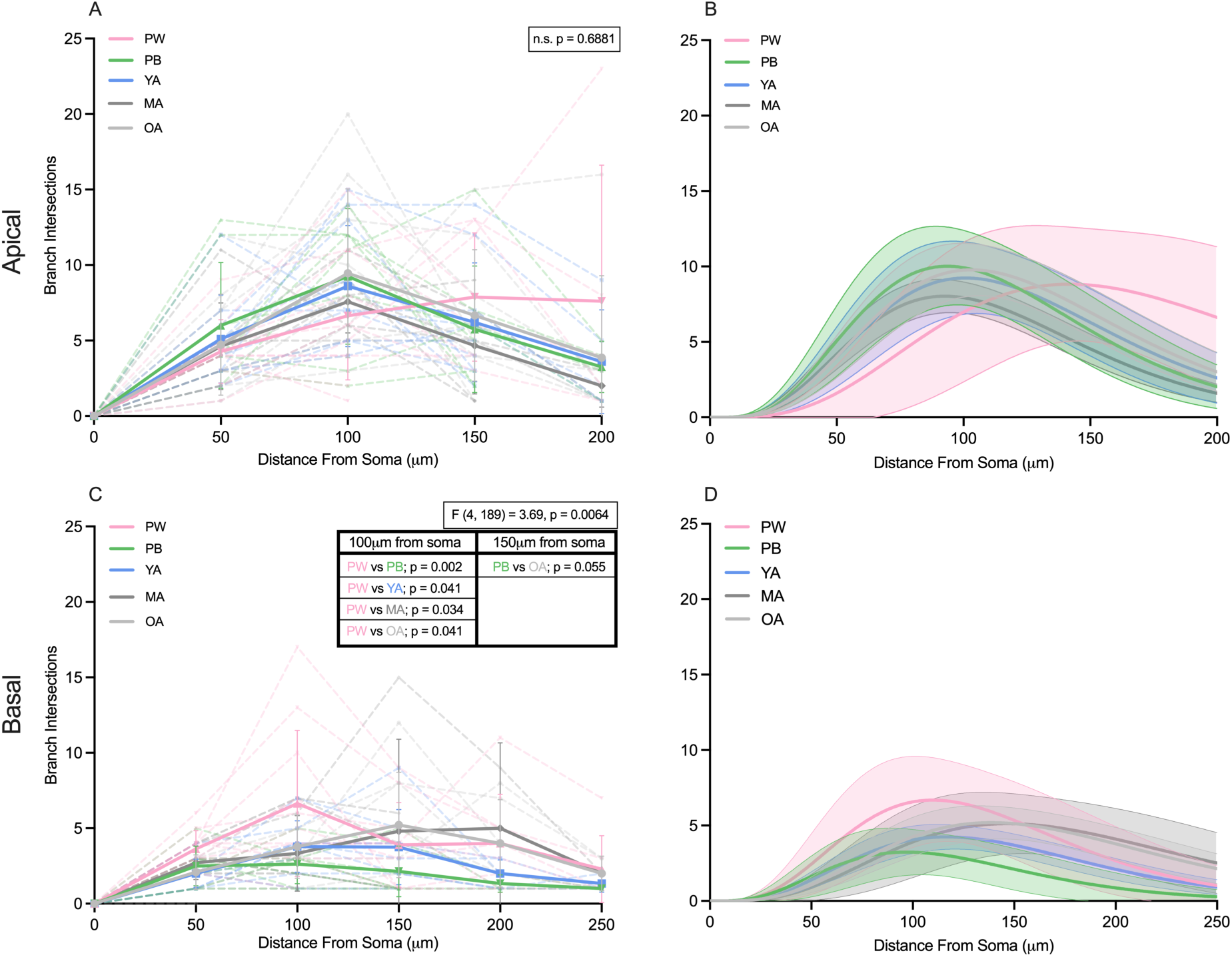
Fusiform cells prune and remodel their basal dendrites with age. *A,C*: Sholl analysis of the apical (*A*) and basal (*C*) dendritic tree. Dashed lines indicate individual neurons, while solid lines indicate means for each age group. Error bars are denoted as standard deviation. *B,D*: Gamma fit curves of each age group for the apical (*B*) and basal (*D*) dendritic tree. Solid lines denote gamma fit to the mean, while shaded regions denote 95% confidence interval bands. 2-way ANOVA (distance and age) used for statistical analysis, with alpha = 0.05. All statistical analysis was conducted on raw data set (*A* and *C*). If significance is found with the omnibus test, Tukey’s multiple comparisons post-hoc test is completed. Statistics located in black boxes denote omnibus calculations. n.s. = not significant. Age groups: PW: preweaning, PB: pubescent, YA: young adult, MA: mature adult, OA: old adult.

## DISCUSSION

We conducted a novel study of age-related electrophysiological and morphological properties of murine FCs, the primary output cell of the DCN. Previous murine slice electrophysiological studies in DCN were conducted between P14-28, (31, 50, 57, 58, 64, 91–107) due to the difficulty of recording from older animals. Our ability to successfully record from the DCN in older animals was facilitated by using a modified slice preparation method (71) and the transstrial cutting angle. We found that there are persistent, slow developing age-related changes in the shapes of action potentials and firing rates of FCs. We also observed remodeling of FC dendritic trees with age. These observations reveal that after the onset of hearing, there are enduring age-related changes in FC morphology and physiology, some of which coincide with the onset of ARHL.

### The Onset of ARHL in 18-month-old CBA/CaJ mice

Our findings of an ABR threshold increase and amplitude decrease in the oldest age group are consistent with prior reports in CBA/CaJ mice (82, 83). These changes are likely caused by either the loss of auditory nerve fiber contacts with inner hair cells or hair cell degeneration, both hallmarks for the onset of ARHL. Our LSFM observations are in agreement with previous studies of CBA/CaJ mice (6, 81, 83, 108) and other species (109) showing that by 18 months of age there is a loss of hair cells (IHC and OHC) at both the apex and base of the cochlea. Such losses at the receptor level would reduce the number and dynamic range of auditory inputs available to the CN. In the United States, ARHL affects one-third of humans aged 65-74 years (3). Although a direct correlation between human and mouse age in years is difficult, studies estimate that laboratory mice 18-24 months old are equivalent to 56-69 years of age in humans (110). This comparison ranks our oldest age group approximate to the biological onset of human ARHL.

### Limitations and comparisons with previous studies

This study provides morphological and physiological measurements of DCN FCs from P15 through P578; however, there are limitations to the interpretation of the data that should be considered. First, in the Sholl analysis, although the average measures of basal and apical dendrite structures are consistent within individual age groups, the measurements were made on a limited number of reconstructed cells. Further analysis of total branch and terminal points from a larger population of FCs would be desirable, perhaps from full reconstructions using light-sheet microscopy from intact DCN rather than neurons in brain slices. In addition, we did not attempt to assess spine density or formally measure dendrite diameters, both of which might be expected to undergo additional age-related changes. These measures would be better made in conjunction with specific structural markers of synapses or using methods with higher spatial resolution.

Methodological differences, including mouse strain, sample preparation, measurement approaches, and data selection criteria exist between our studies and those from other laboratories, complicating comparisons. While we used approaches similar to our previous studies in the DCN (74, 75), our findings may have quantitative differences compared to data from other labs. These results also depend on our classification of age ranges for mice, which were established at the start of the study and were not further adjusted.

After sealing the pipette to the cell membrane in voltage-clamp mode we measured pipette capacitance and then applied 50% compensation of this pipette capacitance in subsequent current clamp recordings, a technique applied in cortex (76). While full compensation of the pipette capacitance results in even faster action potential kinetics, in our hands this can introduce ringing in the action potential waveform, reflecting excessive gain and phase shifts at higher frequencies. To minimize these alterations, we low-pass filtered at 20 kHz to capture the shapes of action potentials with less distortion. While our methods are well-defined, consistently applied across ages, and reproducible, these data will be different than those obtained in other laboratories utilizing different compensation parameters.

### Fusiform Cell Excitability

The intrinsic excitability of neurons in mice develops rapidly during the first 2 weeks of life with continuing, but slower, development thereafter (67, 68, 111). However, intrinsic excitability is not a static phenomenon and can undergo homeostatic changes in response to changes in synaptic input and overall spiking levels (112), including in the CN (113, 114). Benites et al. 2023 (64) showed an increase in spontaneous spiking activity between P12 and P22, accompanied by an increase in the rate of rise and peak action potential amplitude, a decrease in the action potential half-width, and a decrease in current required to elicit spiking. With the development of ARHL, it would be expected that sensory-driven activity in the auditory nerve will decrease, and the most likely homeostatic change in neurons targeted by the nerve is an increase in intrinsic excitability.

The most pronounced differences that we observed were in the action potential shape and maximal firing rate, each appearing to have a monotonic age-dependent shift between the preweaning and old adult ages. The action potential shape differences were primarily driven by faster repolarization slopes, likely representing an increased conductance though fast K^+^ channels. The decreases in the action potential width with age may contribute to the slightly (but significantly) faster firing rates for large current injections in multiple ways. First, the faster-falling slope and narrower action potential width could decrease accumulated Na^+^ channel inactivation. Second, decreased activation of Ca^2+^ channels could reduce activation of SK and/or BK channels, reducing activation of slower K^+^ channels and shortening the medium-AHP. Third, additional slower K^+^ channels that enhance slower AHPs and raise AP threshold are less activated during briefer APs (27, 111). A similar pattern of age-related excitability changes have been reported in other areas of the brain. Chang et al. (115) found that action potentials in older monkey prefrontal cortex were smaller and narrower, with subsequent increase in firing rates. Similarly, layer 2/3 pyramidal cells in the human cerebral cortex become narrower and smaller in late adulthood, although an increase in firing rate was less clear (116). In mouse somatosensory cortex, firing rates were increased in both adapting and non-adapting layer 5 neurons t18-29 months compared to 2-6 months(117). Our work suggests that these increases in firing rate and changes in action potential shape reported in cortex may generalize to aging brainstem nuclei.

Age-related changes in spike shape can have significant consequences on the expression of synaptic plasticity in the DCN. Mouse DCN FCs show a Hebbian spike-timing-dependent plasticity at their parallel fiber inputs (106, 118). Similar plasticity has been reported *in vivo* for interactions between somatosensory and auditory stimuli (119, 120). In general, timing-dependent synaptic plasticity depends on the coincidence of synaptic inputs with Ca^2+^ influx into dendrites from back-propagating action potentials, and dendritic back-propagating action potentials that open calcium channels are known to occur in both FCs and cartwheel cells (121, 122). Such backpropagating action potentials can also provide depolarization to open NMDA receptors that are also present in these cells (50). The shape of action potentials greatly affect the time course of Ca^2+^ channel opening and the voltage-dependent unblocking NMDA receptors. In the absence of other channel compensation, faster action potentials will reduce dendritic action potential amplitude and decrease postsynaptic Ca^2+^ in the dendrites. This reduced Ca^2+^ will likely diminish activation of mechanisms that lead to long-term synaptic plasticity. These observations predict that backpropagating action potential-dependent synaptic plasticity at the parallel-fiber to FC synapse would be depressed in the old mice as compared to young mice.

### Fusiform Cells Exhibit Basal Dendritic Remodeling at the Onset of ARHL

We observed a change in basal dendritic branching between the preweaning age group compared to the middle 3 ages, suggesting that dendritic pruning could be occurring in FCs after the onset of hearing. This early pruning of the basal dendrites is followed by an increase in dendritic branching distal to the soma in the oldest age group (see Figs. 8 and 9). These observations raise the question: Are these altered morphological features related to changes in the level of auditory afferent inputs associated with age?

The increase in FC firing rates during the onset of ARHL could result from a decrease of low-SR spiral ganglion neuronal innervation of the CN, as previously suggested for bushy cells (123). A possible compensatory mechanism for FCs affected by the loss of SGN innervation could be morphological changes in the basal dendritic structure. It was shown in other cell types that dendritic changes occur due to aging (124, 125). We observed a decrease in basal dendritic complexity branching at 100 µm from the FC soma in the preweaning age group compared to all other age groups, indicating potential basal dendrite pruning shortly after hearing onset (Fig.9C). Interestingly, basal branching increased at 150 µm from the soma in the old adult age group compared to the pubescent age group, indicating remodeling of the basal dendritic tree (Fig.9C, D). From the gamma function curve fits, we were able to quantitatively identify where peak branching occurred, relative to the soma. Interestingly, aligned with the increase of basal dendritic branching in the old adult mice, we observe peak branching occurring further from the soma with age (Fig.9D, Table 2). These findings indicate that the basal dendritic tree of FCs undergo structural changes in response to altered spiral ganglion neuron inputs.

In contrast to the basal dendrites, the preweaning age group showed maximal branching of the apical dendrites further from the soma than all other age groups (Fig.9B, Table 1). This can be interpreted as a refinement of apical dendrites in fusiform cells with maturation. However, we observed only modest changes in apical dendritic complexity beyond pubescence, suggesting that the slight structural changes in FC morphology are related to inputs from the parallel fibers of aging granule cells (Fig.9A, B). Further studies of DCN granule cell activity in the context of age-related hearing loss would aid in our understanding of the network effects of the DCN with age.

### Future Directions for Age-Related Studies in the CN

It has been suggested that age-related hearing loss diminishes inhibitory function, which in turn can alter sensory responses (14, 36, 37, 126). Within the cochlear nucleus and thalamus decreased inhibitory signaling from age-related hearing loss induced hyperexcitability in excitatory neurons (14, 20, 39). D-stellate cells, an inhibitory cell in the VCN, show decreased firing in response to auditory nerve stimulation associated with age related hearing loss (38), and are implicated in the direct inhibition of FCs (127–129). FCs also receive inhibition locally from tuberculoventral cells and cartwheel cells (57, 59, 60, 62, 130, 131), two cell types that are understudied in the context of age-related hearing loss. Here, we have extended *in vitro* brain slice recordings in the DCN to much older animals than previously reported, opening opportunities for further studies to gain insights into the physiological changes of the aging auditory system and accompanying age-related hearing loss. Thus, an important future direction would be to clarify the physiological and morphological properties of these local inhibitory cells with age and elucidate their contribution to the enhanced firing of FCs associated with age-related hearing loss.

## DATA AVAILABILITY

Source data (analyzed measurements) is available at FigShare: 10.6084/m9.figshare.29508389.

## GRANTS

This work was supported by the NIDCD R01 DC019053 (PBM) and NINDS Ruth L. Kirschstein Predoctoral Individual NRSA F31 NS129291-01A1 (RJE). Confocal microscopy and Imaris analysis at the UNC Neuroscience Microscopy Core (RRID:SCR_019060), supported, in part, by funding from the NIH-NINDS Neuroscience Center Support Grant P30 NS045892 and the NIH-NICHD Intellectual and Developmental Disabilities Research Center Support Grant P50HD103573. Light sheet fluorescence microscopy conducted at the Microscopy Services Laboratory, Department of Pathology and Laboratory Medicine, is supported in part by P30 CA016086 Cancer Center Core Support Grant to the UNC Lineberger Comprehensive Cancer Center. Research reported in this publication was supported in part by the North Carolina Biotech Center Institutional Support Grant 2016-IDG-1016. These funding agencies had no role in study design, data collection and interpretation, or decision to present this work.

## DISCLOSURES

The authors have no disclosures to report.

## DISCLAIMERS

The authors have no disclaimers.

## AUTHOR CONTRIBUTIONS

R.J.E, M.R.K., and P.B.M. conceptualized the research; R.J.E, M.R.K., K.A.H., and M.P.L. performed experimentation; R.J.E., and P.B.M. analyzed the data; R.J.E., M.R.K., K.A.H., and P.B.M. interpreted the results; R.J.E., K.A.H., and P.B.M. prepared figures; R.J.E drafted the initial manuscript; R.J.E., M.R.K., K.A.H., and P.B.M edited and revised the manuscript; R.J.E., M.R.K., K.A.H., M.P.L., and P.B.M. approved the final version of manuscript.

## SUPPLEMENTAL MATERIAL

**Table S1:** Passive Properties.

**SUPPLEMENTAL MATERIAL**718 **Table S1: Passive Properties**
| Age category |  |  |  |  |  | F<sup>2</sup> | D.F. | p-value<sup>2</sup> |
| --- | --- | --- | --- | --- | --- | --- | --- | --- |
| Measure | Prewaning<br>N = 22 | Pubescent<br>N = 48 | Young Adult<br>N = 26 | Mature Adult<br>N = 40 | Old Adult<br>N = 27 |  |  |  |
| RMP (mV) | -69.7<br>(3.5, 22)<sup>1</sup> | -69.0<br>(5.2, 48) | -72.3<br>(5.8, 26) | -69.6<br>(5.1, 40) | -69.4<br>(3.4, 27) | 2.20<br>4 | (4, 158) | 0.0<br>71 |
| R<sub>in</sub> (MΩm) | 58.7<br>(23.2, 21) | 48.0<br>(17.9, 44) | 54.6<br>(20.8, 26) | 47.3<br>(13.3, 39) | 55.0<br>(23.0, 27) | 1.95<br>6 | (4, 152) | 0.1<br>04 |
| t<sub>m</sub> (ms) | 11.2<br>(4.0, 21) | 10.0<br>(3.0, 44) | 11.2<br>(5.8, 26) | 10.5<br>(3.4, 39) | 10.5<br>(5.8, 27) | 0.42<br>8 | (4, 152) | 0.7<br>88 |

<sup>1</sup>Mean (SD, N)<sup>2</sup>One-way analysis of means (not assuming equal variances)719 **Table S2: Spike Shape**

**Table S2:**
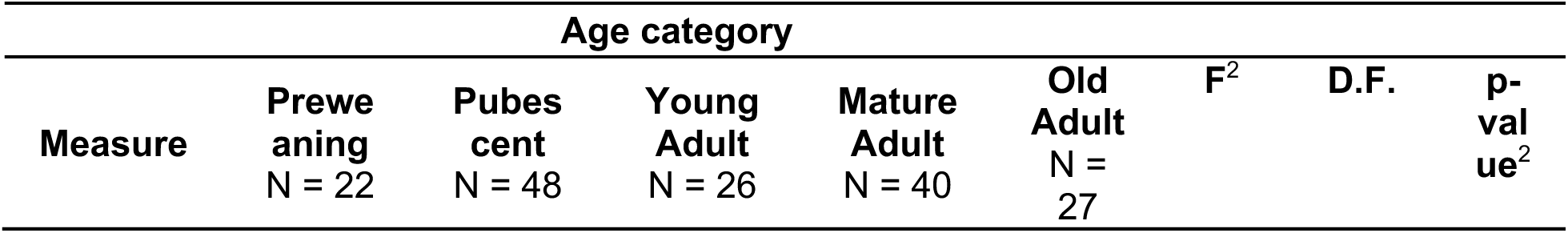
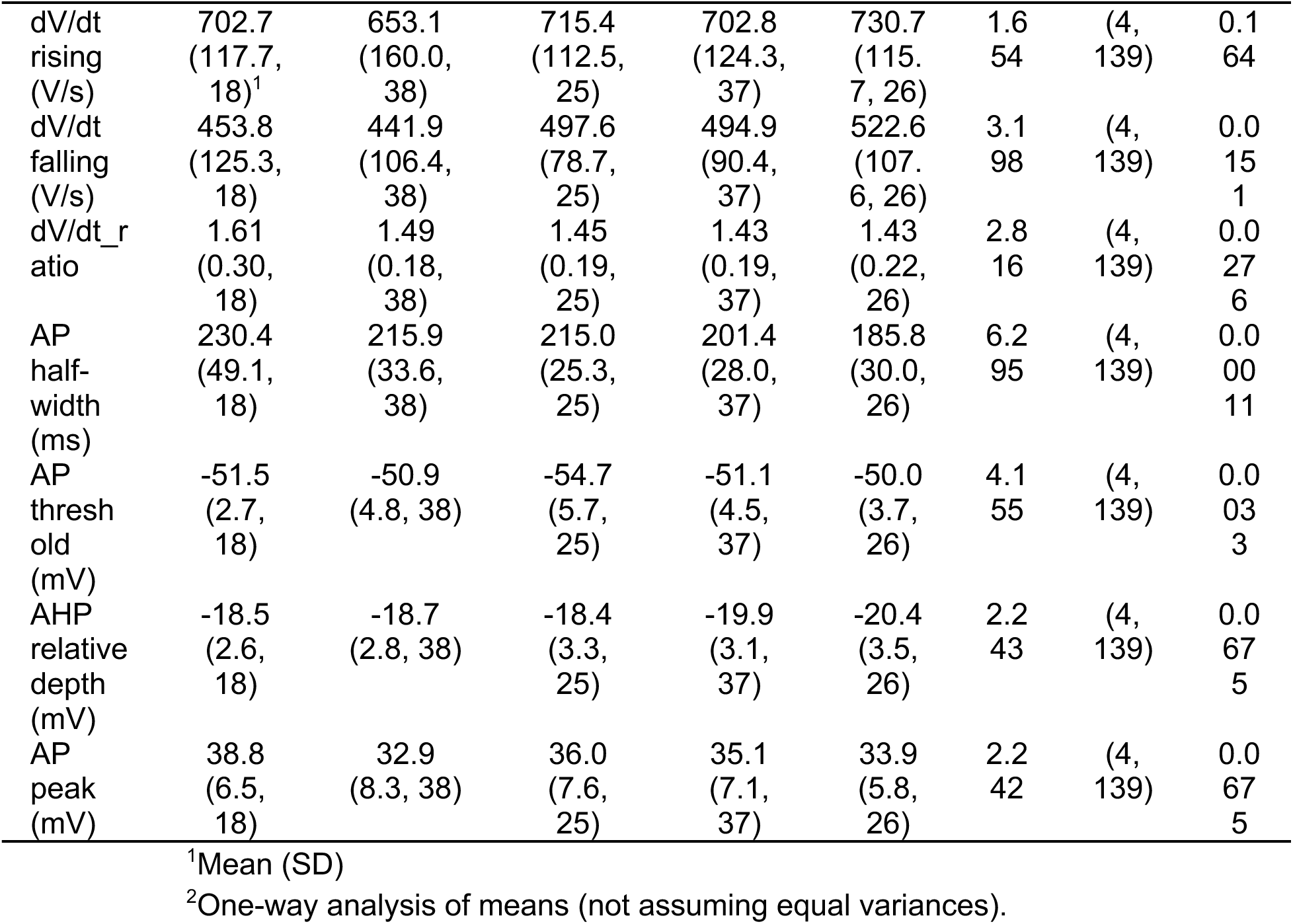
Spike Shape.

**Table S3:**
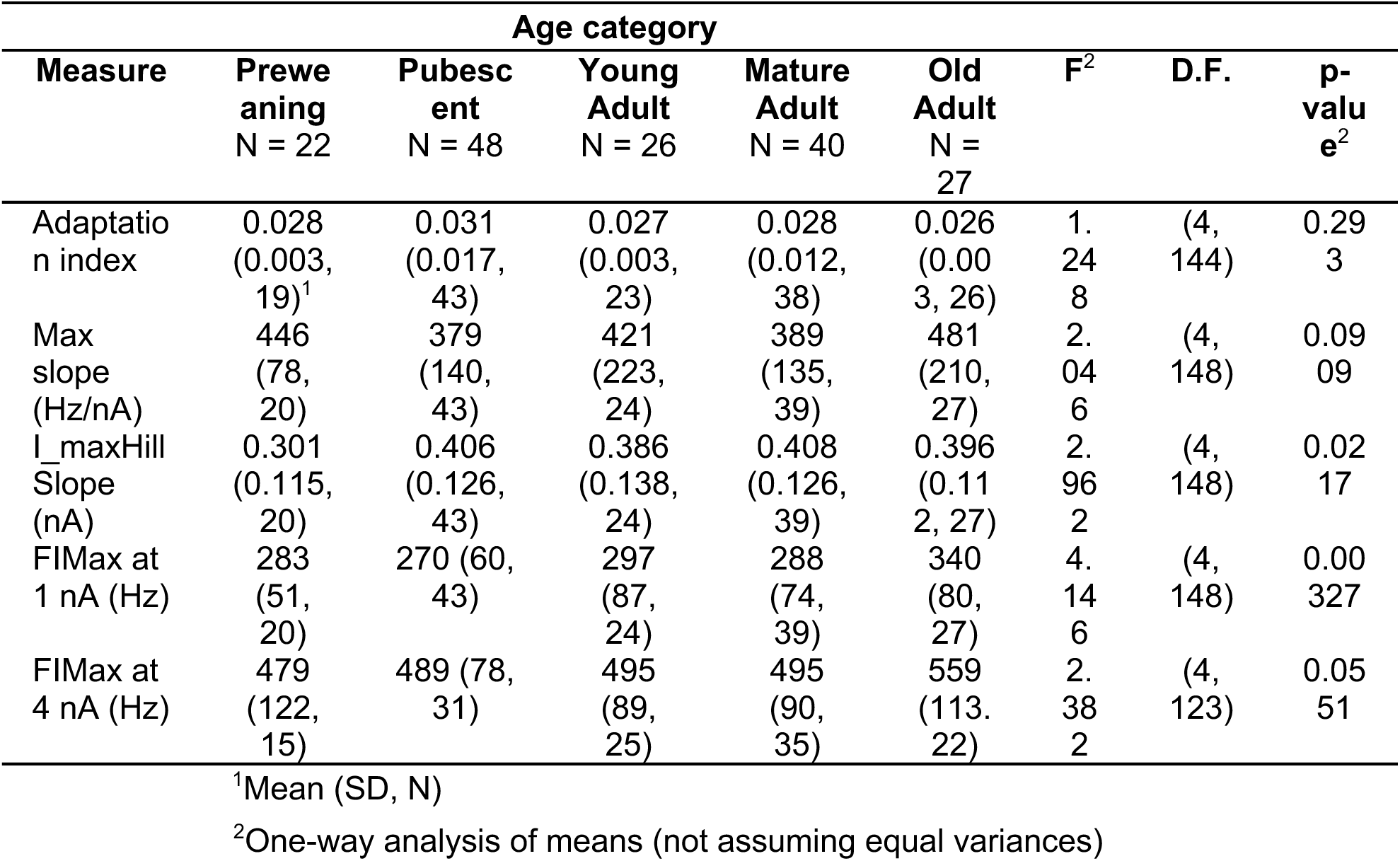
Firing Rates.

**Table S4:**
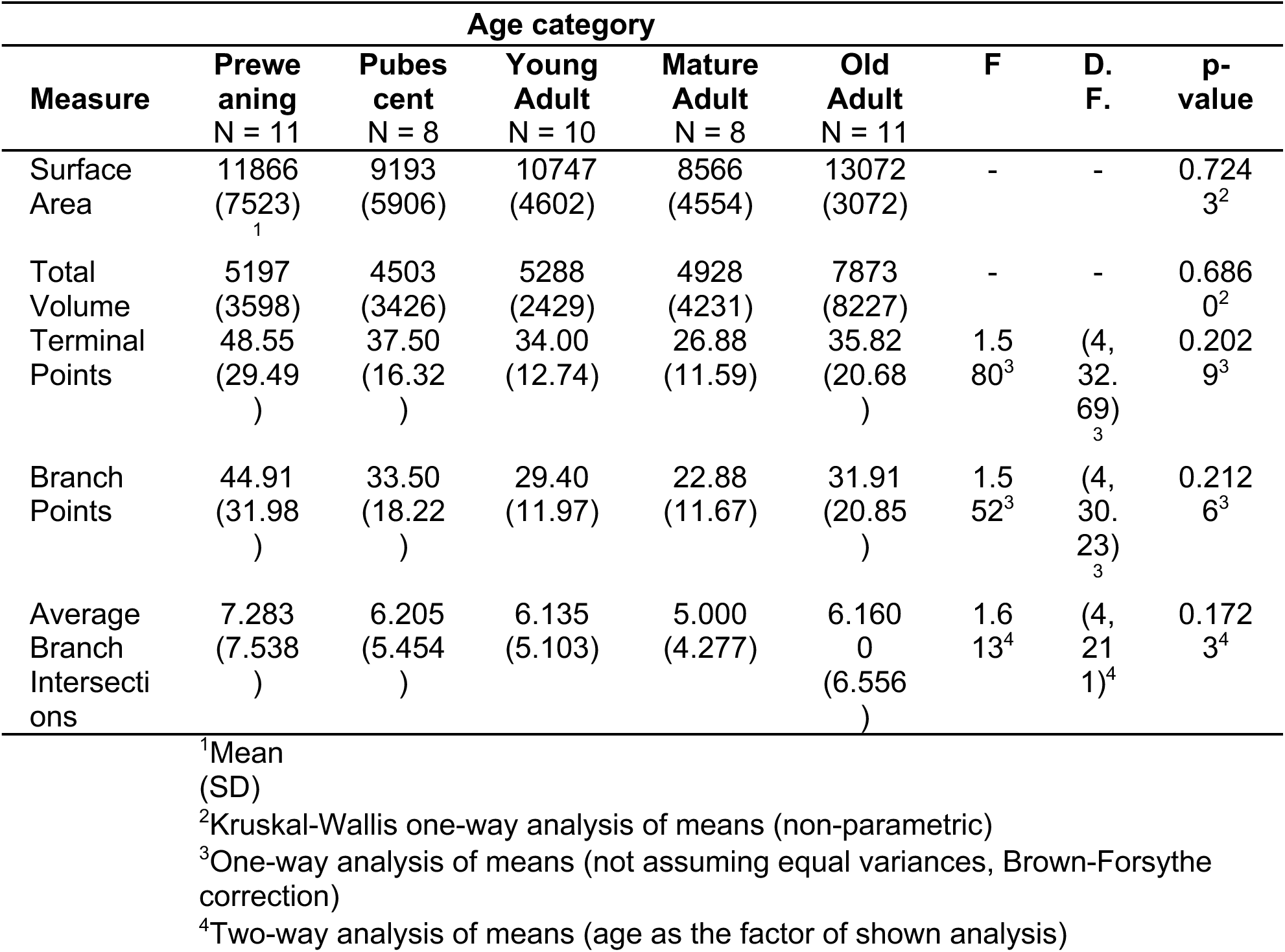
Whole Cell Morphology.

**Table S5:**
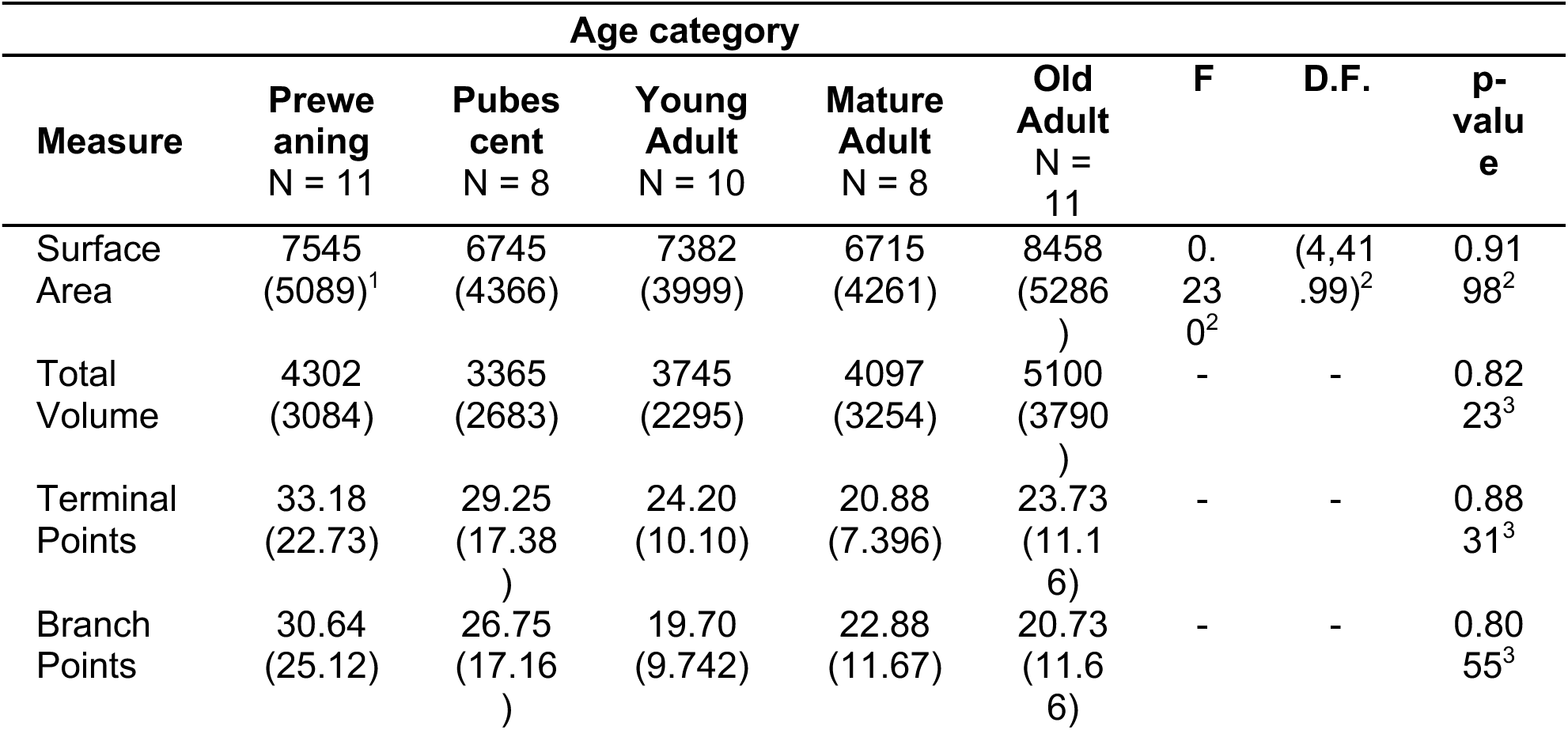
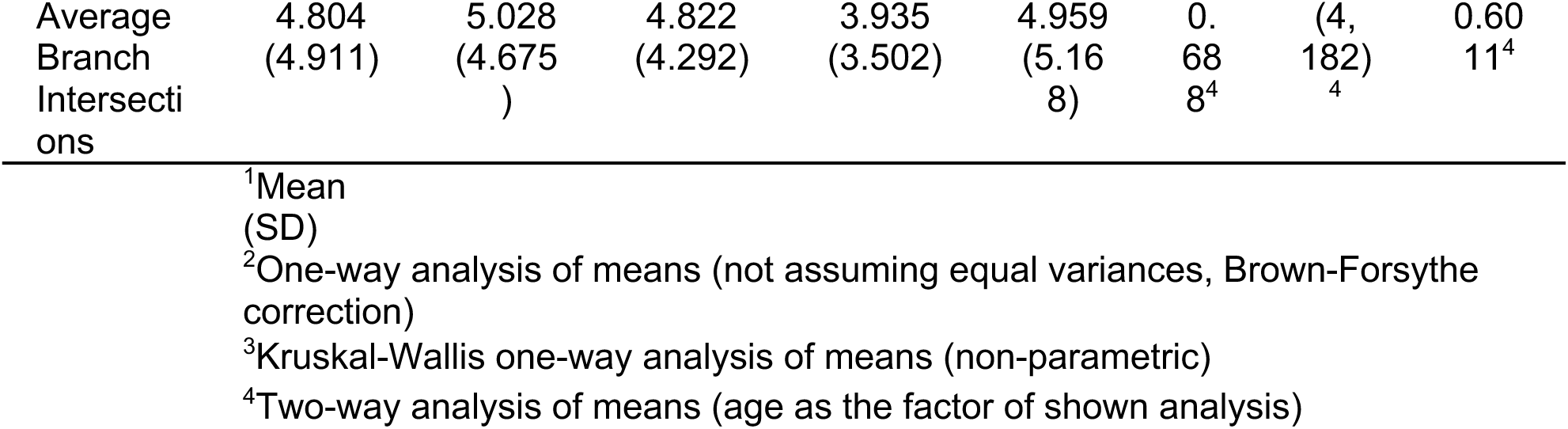
Apical Dendrite Morphology.

**Table S5:**
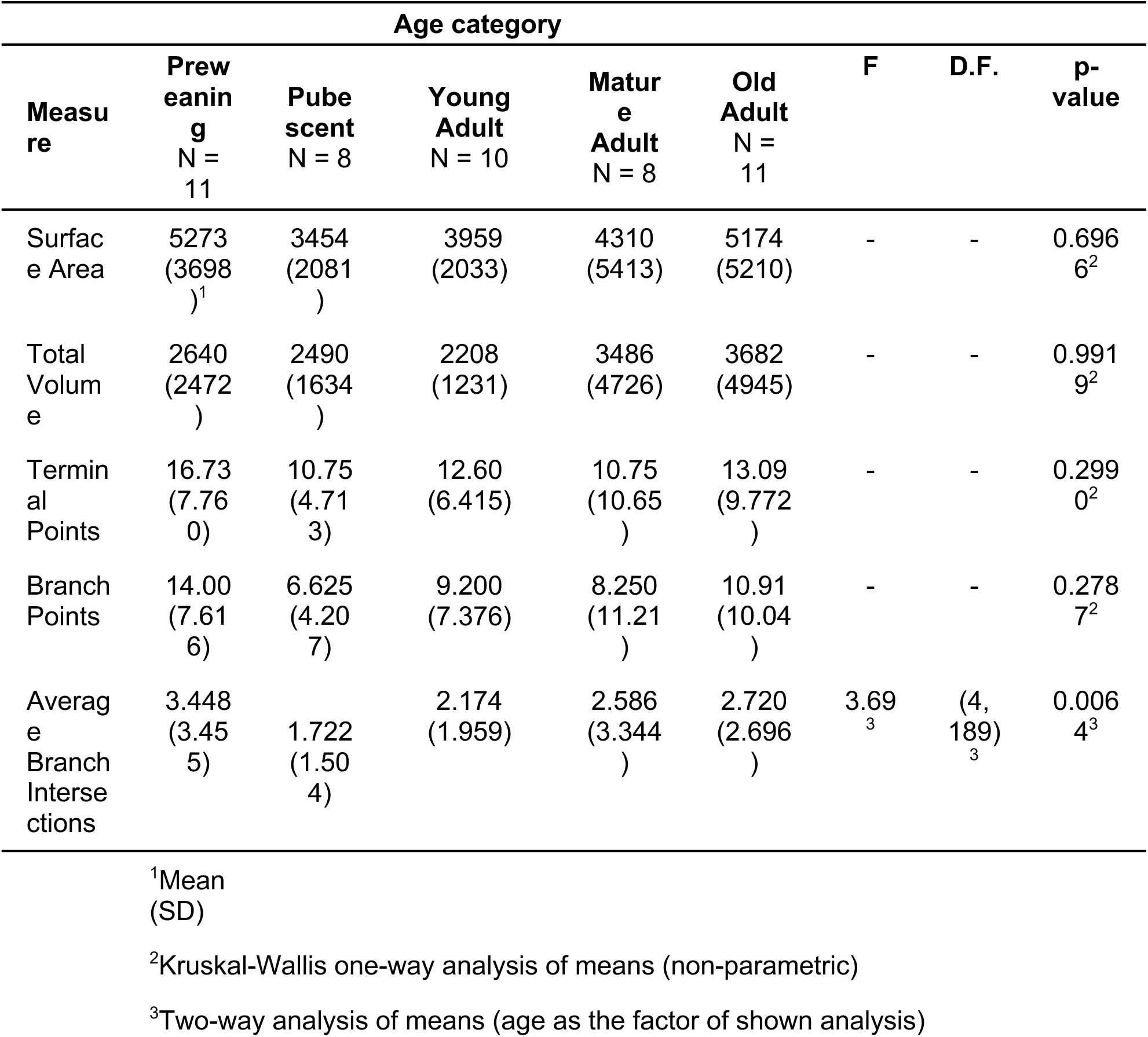
Basal Dendrite Morphology.

| Measure | Age category |  |  |  |  | F | D.F. | p-value |
| --- | --- | --- | --- | --- | --- | --- | --- | --- |
|  | Prew<br>eaning<br>N = 11 | Pube<br>scent<br>N = 8 | Young<br>Adult<br>N = 10 | Matur<br>e<br>Adult<br>N = 8 | Old<br>Adult<br>N = 11 |  |  |  |
| Surface Area | 5273<br>(3698)<sup>1</sup> | 3454<br>(2081) | 3959<br>(2033) | 4310<br>(5413) | 5174<br>(5210) | - | - | 0.6966<sup>2</sup> |
| Total Volume | 2640<br>(2472) | 2490<br>(1634) | 2208<br>(1231) | 3486<br>(4726) | 3682<br>(4945) | - | - | 0.9919<sup>2</sup> |
| Terminal Points | 16.73<br>(7.760) | 10.75<br>(4.713) | 12.60<br>(6.415) | 10.75<br>(10.65) | 13.09<br>(9.772) | - | - | 0.2990<sup>2</sup> |
| Branch Points | 14.00<br>(7.616) | 6.625<br>(4.207) | 9.200<br>(7.376) | 8.250<br>(11.21) | 10.91<br>(10.04) | - | - | 0.2787<sup>2</sup> |
| Average Branch Intersections | 3.448<br>(3.455) | 1.722<br>(1.504) | 2.174<br>(1.959) | 2.586<br>(3.344) | 2.720<br>(2.696) | 3.69<sup>3</sup> | (4, 189)<sup>3</sup> | 0.0064<sup>3</sup> |

<sup>1</sup>Mean (SD)
<sup>2</sup>Kruskal-Wallis one-way analysis of means (non-parametric)
<sup>3</sup>Two-way analysis of means (age as the factor of shown analysis)

**Table S6:**
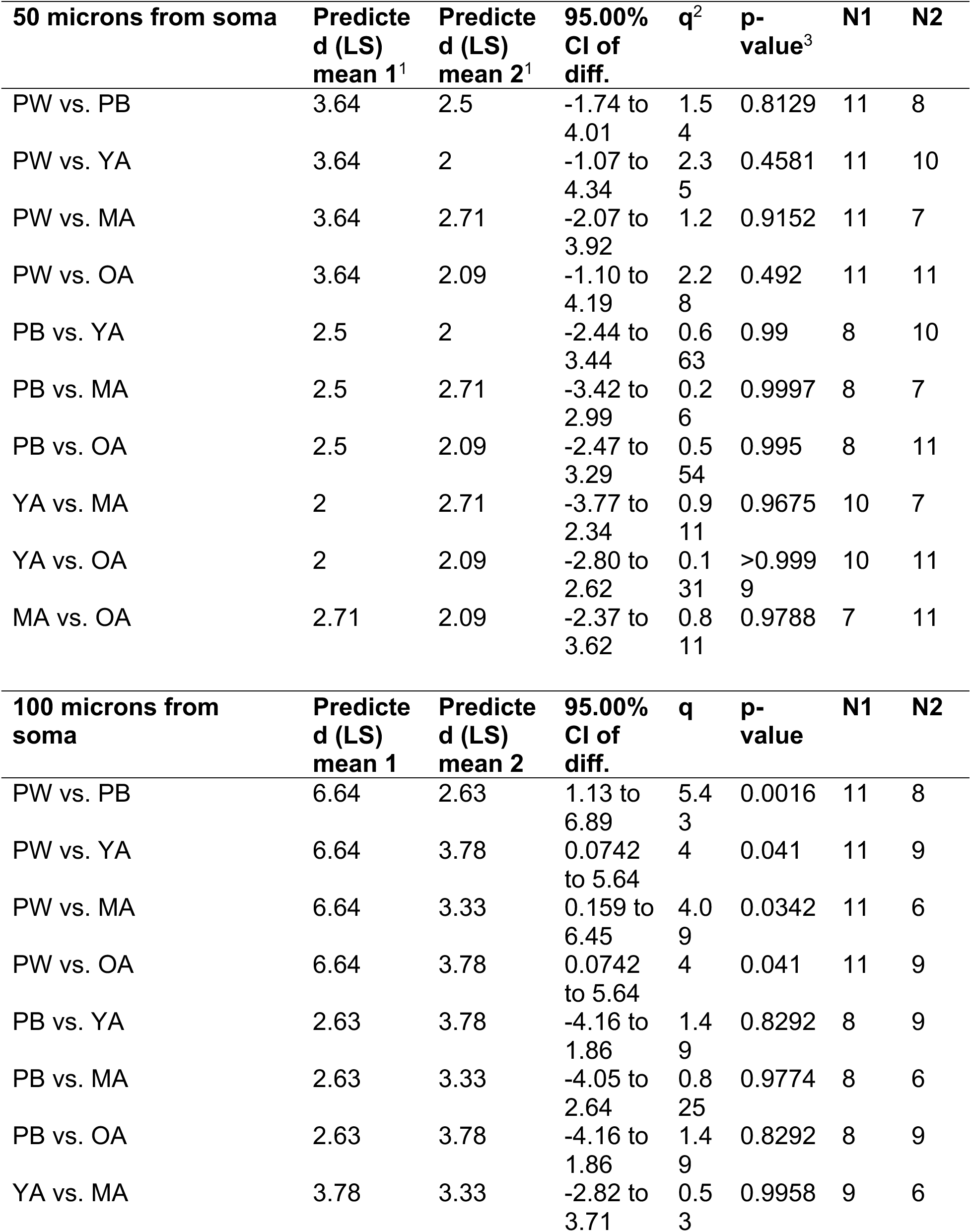
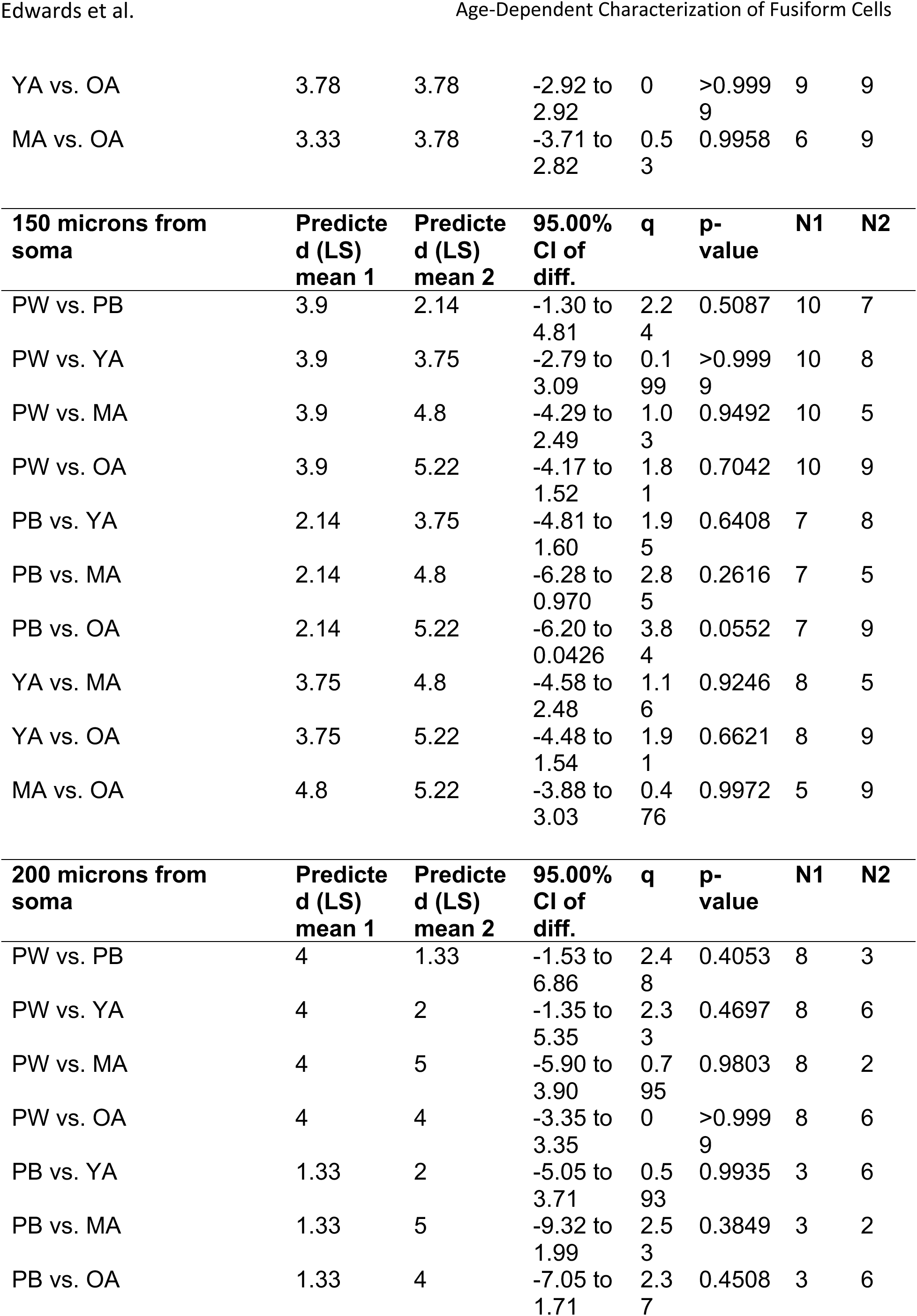
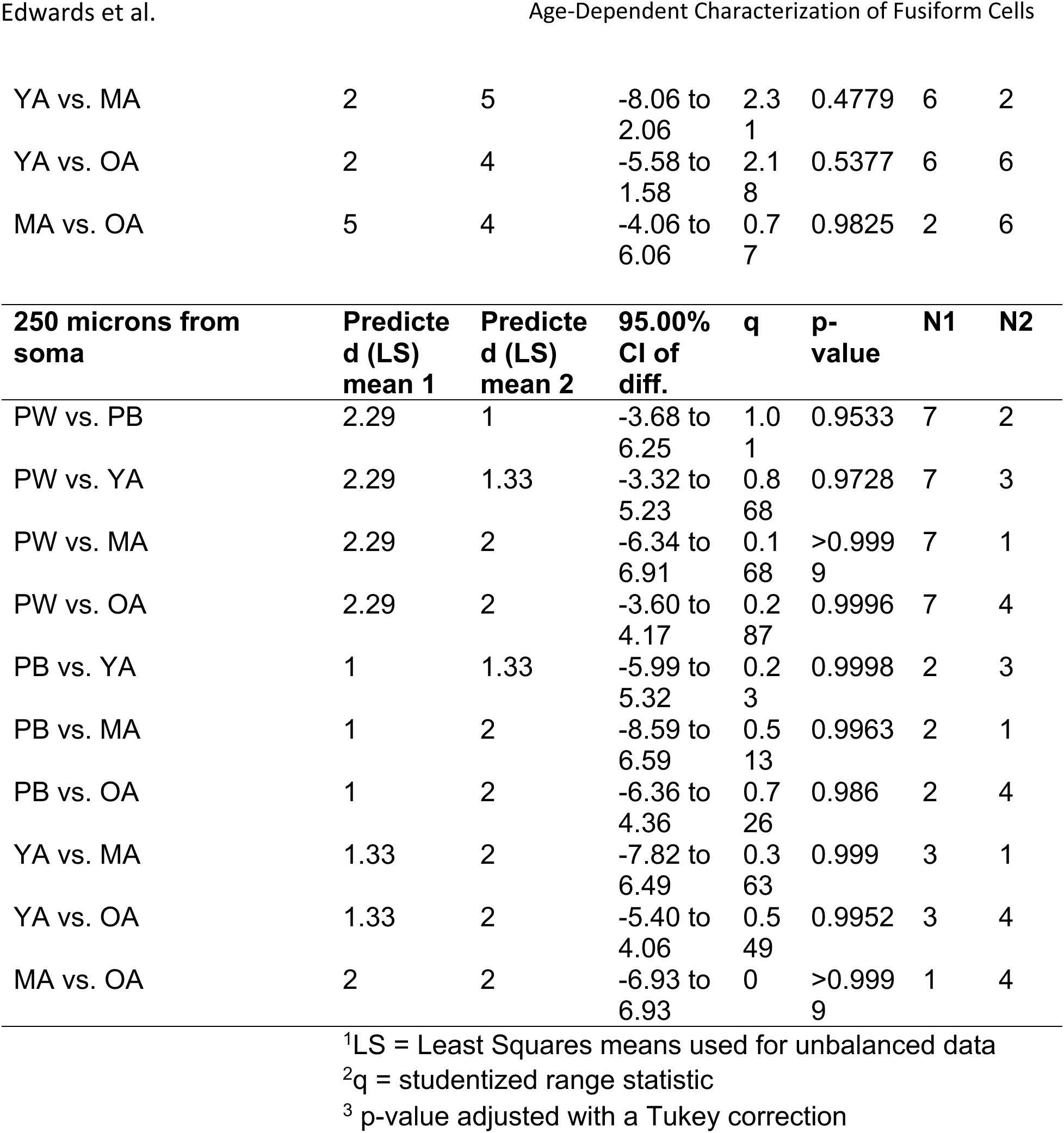
Basal Dendrite Multiple Comparisons Post-hoc Test.

## Notes

### Competing Interest Statement

The authors have declared no competing interest.

### Funding

- National Institutes of Health: R01 DC 019053, F31 NS129291-01

