## Supplemental Tables and Figure for "Fusiform Cells in the Dorsal Cochlear Nucleus Change Intrinsic Electrophysiological Properties and Morphologically Remodel Their Basal Dendrites with Age"

**Supplemental Figure 1**

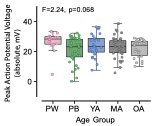

Figure S1: The absolute peak voltage of action potentials does not change with age. These are the same measurements shown in Figure 4I, but not adjusted for the action potential threshold. PW: preweaning; PB: pubescent; YA: young adult; MA: mature adult; OA: old adult. See main text for age ranges. Statistical analysis: 1-way ANOVA.

Supplemental Tables:

Tables S1, S2 and S3 list the means, standard deviations, and sample sizes of the measured values in each age group. The N values in the top row indicate the total number of cells associated with each group in the study. The right-most columns show the results of a 1-way ANOVA. The degrees of freedom for the ANOVAs vary between measurements according to the whether appropriate data was collected, and whether that data was usable for a particular analysis. For example, in the Preweaning and Mature adult groups, one cell did not have a protocol with a small hyperpolarizing step that could be used to measure Rin and the membrane time constant; in the Pubescent age group there were 4 cells lacking this data.

Table S1: Passive Properties

|  | **Age category** | | | | |  |  | |
| --- | --- | --- | --- | --- | --- | --- | --- | --- |
| **Measure** | **Preweaning**  N = 22 | **Pubescent**  N = 48 | **Young Adult**  N = 26 | **Mature Adult**  N = 40 | **Old Adult**  N = 27 | **F**^2^ | **D.F.** | **p-value**^2^ |
| RMP (mV) | -69.7 (3.5, 22)^1^ | -69.0 (5.2, 48) | -72.3 (5.8, 26) | -69.6 (5.1, 40) | -69.4 (3.4, 27) | 2.204 | (4, 158) | 0.071 |
| R_in_ (MOhm) | 58.7 (23.2, 21) | 48.0 (17.9, 44) | 54.6 (20.8, 26) | 47.3 (13.3, 39) | 55.0 (23.0, 27) | 1.956 | (4, 152) | 0.104 |
| τ_m_ (ms) | 11.2 (4.0, 21) | 10.0 (3.0, 44) | 11.2 (5.8, 26) | 10.5 (3.4, 39) | 10.5 (5.8, 27) | 0.428 | (4, 152) | 0.788 |
|  | ^1^Mean (SD, N) | | | | | | | |
|  | ^2^One-way analysis of means (not assuming equal variances) | | | | | | | |

Table S2: Spike Shape

|  | **Age category** | | | | |  |  | |
| --- | --- | --- | --- | --- | --- | --- | --- | --- |
| **Measure** | **Preweaning**  N = 22 | **Pubescent**  N = 48 | **Young Adult**  N = 26 | **Mature Adult**  N = 40 | **Old Adult**  N = 27 | **F**^2^ | **D.F.** | **p-value**^2^ |
| dV/dt rising (V/s) | 702.7 (117.7, 18)^1^ | 653.1 (160.0, 38) | 715.4 (112.5, 25) | 702.8 (124.3, 37) | 730.7 (115.7, 26) | 1.654 | (4, 139) | 0.164 |
| dV/dt falling (V/s) | 453.8 (125.3, 18) | 441.9 (106.4, 38) | 497.6 (78.7, 25) | 494.9 (90.4, 37) | 522.6 (107.6, 26) | 3.198 | (4, 139) | 0.0151 |
| dV/dt_ratio | 1.61 (0.30, 18) | 1.49 (0.18, 38) | 1.45 (0.19, 25) | 1.43 (0.19, 37) | 1.43 (0.22, 26) | 2.816 | (4, 139) | 0.0276 |
| AP half-width (ms) | 230.4 (49.1, 18) | 215.9 (33.6, 38) | 215.0 (25.3, 25) | 201.4 (28.0, 37) | 185.8 (30.0, 26) | 6.295 | (4, 139) | 0.00011 |
| AP threshold (mV) | -51.5 (2.7, 18) | -50.9 (4.8, 38) | -54.7 (5.7, 25) | -51.1 (4.5, 37) | -50.0 (3.7, 26) | 4.155 | (4, 139) | 0.0033 |
| AHP relative depth (mV) | -18.5 (2.6, 18) | -18.7 (2.8, 38) | -18.4 (3.3, 25) | -19.9 (3.1, 37) | -20.4 (3.5, 26) | 2.243 | (4, 139) | 0.0675 |
| AP peak (mV) | 38.8 (6.5, 18) | 32.9 (8.3, 38) | 36.0 (7.6, 25) | 35.1 (7.1, 37) | 33.9 (5.8, 26) | 2.242 | (4, 139) | 0.0675 |
|  | ^1^Mean (SD) | | | | | | | |
|  | ^2^One-way analysis of means (not assuming equal variances). | | | | | | | |

Table S3: Firing Rates

|  | **Age category** | | | | |  |  | |
| --- | --- | --- | --- | --- | --- | --- | --- | --- |
| **Measure** | **Preweaning**  N = 22 | **Pubescent**  N = 48 | **Young Adult**  N = 26 | **Mature Adult**  N = 40 | **Old Adult**  N = 27 | **F**^2^ | **D.F.** | **p-value**^2^ |
| Adaptation index | 0.028 (0.003, 19)^1^ | 0.031 (0.017, 43) | 0.027 (0.003, 23) | 0.028 (0.012, 38) | 0.026 (0.003, 26) | 1.248 | (4, 144) | 0.293 |
| Max slope (Hz/nA) | 446 (78, 20) | 379 (140, 43) | 421 (223, 24) | 389 (135, 39) | 481 (210, 27) | 2.046 | (4, 148) | 0.0909 |
| I_maxHillSlope (nA) | 0.301 (0.115, 20) | 0.406 (0.126, 43) | 0.386 (0.138, 24) | 0.408 (0.126, 39) | 0.396 (0.112, 27) | 2.962 | (4, 148) | 0.0217 |
| FIMax at 1 nA (Hz) | 283 (51, 20) | 270 (60, 43) | 297 (87, 24) | 288 (74, 39) | 340 (80, 27) | 4.146 | (4, 148) | 0.00327 |
| FIMax at 4 nA (Hz) | 479 (122, 15) | 489 (78, 31) | 495 (89, 25) | 495 (90, 35) | 559 (113. 22) | 2.382 | (4, 123) | 0.0551 |
|  | ^1^Mean (SD, N) | | | | | | | |
|  | ^2^One-way analysis of means (not assuming equal variances) | | | | | | | |

Table S4: Whole Cell Morphology

|  | **Age category** | | | | | | |  |  |  |
| --- | --- | --- | --- | --- | --- | --- | --- | --- | --- | --- |
| **Measure** | **Preweaning**  N = 11 | **Pubescent**  N = 8 | | **Young Adult**  N = 10 | | **Mature Adult**  N = 8 | **Old Adult**  N = 11 | **F** | **D.F.** | **p-value** |
| Surface Area | 11866 (7523)^1^ | 9193 (5906) | | 10747 (4602) | | 8566 (4554) | 13072 (3072) | - | - | 0.7243^2^ |
| Total Volume | 5197 (3598) | 4503 (3426) | | 5288 (2429) | | 4928 (4231) | 7873 (8227) | - | - | 0.6860^2^ |
| Terminal Points | 48.55 (29.49) | 37.50 (16.32) | | 34.00 (12.74) | | 26.88 (11.59) | 35.82 (20.68) | 1.580^3^ | (4,32.69)^3^ | 0.2029^3^ |
| Branch Points | 44.91 (31.98) | 33.50 (18.22) | | 29.40 (11.97) | | 22.88 (11.67) | 31.91 (20.85) | 1.552^3^ | (4,30.23)^3^ | 0.2126^3^ |
| Average Branch Intersections | 7.283 (7.538) | 6.205 (5.454) | | 6.135 (5.103) | | 5.000 (4.277) | 6.1600 (6.556) | 1.613^4^ | (4, 211)^4^ | 0.1723^4^ |
|  | ^1^Mean (SD) |  |  | |  |  |  |  |  |  |
|  | ^2^Kruskal-Wallis one-way analysis of means (non-parametric) | | | | | | | | | |
|  | ^3^One-way analysis of means (not assuming equal variances, Brown-Forsythe correction) | | | | | | | | | |
|  | ^4^Two-way analysis of means (age as the factor of shown analysis) | | | | | | | | | |

Table S5: Apical Dendrite Morphology

|  | **Age category** | | | | | | |  |  |  |
| --- | --- | --- | --- | --- | --- | --- | --- | --- | --- | --- |
| **Measure** | **Preweaning**  N = 11 | **Pubescent**  N = 8 | | **Young Adult**  N = 10 | | **Mature Adult**  N = 8 | **Old Adult**  N = 11 | **F** | **D.F.** | **p-value** |
| Surface Area | 7545 (5089)^1^ | 6745 (4366) | | 7382 (3999) | | 6715 (4261) | 8458 (5286) | 0.230^2^ | (4,41.99)^2^ | 0.9198^2^ |
| Total Volume | 4302 (3084) | 3365 (2683) | | 3745 (2295) | | 4097 (3254) | 5100 (3790) | - | - | 0.8223^3^ |
| Terminal Points | 33.18 (22.73) | 29.25 (17.38) | | 24.20 (10.10) | | 20.88 (7.396) | 23.73 (11.16) | - | - | 0.8831^3^ |
| Branch Points | 30.64 (25.12) | 26.75 (17.16) | | 19.70 (9.742) | | 22.88 (11.67) | 20.73 (11.66) | - | - | 0.8055^3^ |
| Average Branch Intersections | 4.804 (4.911) | 5.028 (4.675) | | 4.822 (4.292) | | 3.935 (3.502) | 4.959 (5.168) | 0.688^4^ | (4, 182)^4^ | 0.6011^4^ |
|  | ^1^Mean (SD) |  |  | |  |  |  |  |  |  |
|  | ^2^One-way analysis of means (not assuming equal variances, Brown-Forsythe correction) | | | | | | | | | |
|  | ^3^Kruskal-Wallis one-way analysis of means (non-parametric) | | | | | | | | | |
|  | ^4^Two-way analysis of means (age as the factor of shown analysis) | | | | | | | | | |

Table S6: Basal Dendrite Morphology

|  | **Age category** | | | | | | |  |  |  |
| --- | --- | --- | --- | --- | --- | --- | --- | --- | --- | --- |
| **Measure** | **Preweaning**  N = 11 | **Pubescent**  N = 8 | | **Young Adult**  N = 10 | | **Mature Adult**  N = 8 | **Old Adult**  N = 11 | **F** | **D.F.** | **p-value** |
| Surface Area | 5273 (3698)^1^ | 3454 (2081) | | 3959 (2033) | | 4310 (5413) | 5174 (5210) | - | - | 0.6966^2^ |
| Total Volume | 2640 (2472) | 2490 (1634) | | 2208 (1231) | | 3486 (4726) | 3682 (4945) | - | - | 0.9919^2^ |
| Terminal Points | 16.73 (7.760) | 10.75 (4.713) | | 12.60 (6.415) | | 10.75 (10.65) | 13.09 (9.772) | - | - | 0.2990^2^ |
| Branch Points | 14.00 (7.616) | 6.625 (4.207) | | 9.200 (7.376) | | 8.250 (11.21) | 10.91 (10.04) | - | - | 0.2787^2^ |
| Average Branch Intersections | 3.448 (3.455) | 1.722 (1.504) | | 2.174 (1.959) | | 2.586 (3.344) | 2.720 (2.696) | 3.69^3^ | (4, 189)^3^ | 0.0064^3^ |
|  | ^1^Mean (SD) |  |  | |  |  |  |  |  |  |
|  | ^2^Kruskal-Wallis one-way analysis of means (non-parametric) | | | | | | | | | |
|  | ^3^Two-way analysis of means (age as the factor of shown analysis) | | | | | | | | | |

Table S7: Basal Dendrite Multiple Comparisons Test

| **50 microns from soma** | **Predicted (LS) mean 1**^1^ | **Predicted (LS) mean 2**^1^ | **95.00% CI of diff.** | **q**^2^ | **p-value**^3^ | **N1** | **N2** |
| --- | --- | --- | --- | --- | --- | --- | --- |
| PW vs. PB | 3.64 | 2.5 | -1.74 to 4.01 | 1.54 | 0.8129 | 11 | 8 |
| PW vs. YA | 3.64 | 2 | -1.07 to 4.34 | 2.35 | 0.4581 | 11 | 10 |
| PW vs. MA | 3.64 | 2.71 | -2.07 to 3.92 | 1.2 | 0.9152 | 11 | 7 |
| PW vs. OA | 3.64 | 2.09 | -1.10 to 4.19 | 2.28 | 0.492 | 11 | 11 |
| PB vs. YA | 2.5 | 2 | -2.44 to 3.44 | 0.663 | 0.99 | 8 | 10 |
| PB vs. MA | 2.5 | 2.71 | -3.42 to 2.99 | 0.26 | 0.9997 | 8 | 7 |
| PB vs. OA | 2.5 | 2.09 | -2.47 to 3.29 | 0.554 | 0.995 | 8 | 11 |
| YA vs. MA | 2 | 2.71 | -3.77 to 2.34 | 0.911 | 0.9675 | 10 | 7 |
| YA vs. OA | 2 | 2.09 | -2.80 to 2.62 | 0.131 | >0.9999 | 10 | 11 |
| MA vs. OA | 2.71 | 2.09 | -2.37 to 3.62 | 0.811 | 0.9788 | 7 | 11 |
| **100 microns from soma** | **Predicted (LS) mean 1** | **Predicted (LS) mean 2** | **95.00% CI of diff.** | **q** | **p-value** | **N1** | **N2** |
| PW vs. PB | 6.64 | 2.63 | 1.13 to 6.89 | 5.43 | 0.0016 | 11 | 8 |
| PW vs. YA | 6.64 | 3.78 | 0.0742 to 5.64 | 4 | 0.041 | 11 | 9 |
| PW vs. MA | 6.64 | 3.33 | 0.159 to 6.45 | 4.09 | 0.0342 | 11 | 6 |
| PW vs. OA | 6.64 | 3.78 | 0.0742 to 5.64 | 4 | 0.041 | 11 | 9 |
| PB vs. YA | 2.63 | 3.78 | -4.16 to 1.86 | 1.49 | 0.8292 | 8 | 9 |
| PB vs. MA | 2.63 | 3.33 | -4.05 to 2.64 | 0.825 | 0.9774 | 8 | 6 |
| PB vs. OA | 2.63 | 3.78 | -4.16 to 1.86 | 1.49 | 0.8292 | 8 | 9 |
| YA vs. MA | 3.78 | 3.33 | -2.82 to 3.71 | 0.53 | 0.9958 | 9 | 6 |
| YA vs. OA | 3.78 | 3.78 | -2.92 to 2.92 | 0 | >0.9999 | 9 | 9 |
| MA vs. OA | 3.33 | 3.78 | -3.71 to 2.82 | 0.53 | 0.9958 | 6 | 9 |
| **150 microns from soma** | **Predicted (LS) mean 1** | **Predicted (LS) mean 2** | **95.00% CI of diff.** | **q** | **p-value** | **N1** | **N2** |
| PW vs. PB | 3.9 | 2.14 | -1.30 to 4.81 | 2.24 | 0.5087 | 10 | 7 |
| PW vs. YA | 3.9 | 3.75 | -2.79 to 3.09 | 0.199 | >0.9999 | 10 | 8 |
| PW vs. MA | 3.9 | 4.8 | -4.29 to 2.49 | 1.03 | 0.9492 | 10 | 5 |
| PW vs. OA | 3.9 | 5.22 | -4.17 to 1.52 | 1.81 | 0.7042 | 10 | 9 |
| PB vs. YA | 2.14 | 3.75 | -4.81 to 1.60 | 1.95 | 0.6408 | 7 | 8 |
| PB vs. MA | 2.14 | 4.8 | -6.28 to 0.970 | 2.85 | 0.2616 | 7 | 5 |
| PB vs. OA | 2.14 | 5.22 | -6.20 to 0.0426 | 3.84 | 0.0552 | 7 | 9 |
| YA vs. MA | 3.75 | 4.8 | -4.58 to 2.48 | 1.16 | 0.9246 | 8 | 5 |
| YA vs. OA | 3.75 | 5.22 | -4.48 to 1.54 | 1.91 | 0.6621 | 8 | 9 |
| MA vs. OA | 4.8 | 5.22 | -3.88 to 3.03 | 0.476 | 0.9972 | 5 | 9 |
| **200 microns from soma** | **Predicted (LS) mean 1** | **Predicted (LS) mean 2** | **95.00% CI of diff.** | **q** | **p-value** | **N1** | **N2** |
| PW vs. PB | 4 | 1.33 | -1.53 to 6.86 | 2.48 | 0.4053 | 8 | 3 |
| PW vs. YA | 4 | 2 | -1.35 to 5.35 | 2.33 | 0.4697 | 8 | 6 |
| PW vs. MA | 4 | 5 | -5.90 to 3.90 | 0.795 | 0.9803 | 8 | 2 |
| PW vs. OA | 4 | 4 | -3.35 to 3.35 | 0 | >0.9999 | 8 | 6 |
| PB vs. YA | 1.33 | 2 | -5.05 to 3.71 | 0.593 | 0.9935 | 3 | 6 |
| PB vs. MA | 1.33 | 5 | -9.32 to 1.99 | 2.53 | 0.3849 | 3 | 2 |
| PB vs. OA | 1.33 | 4 | -7.05 to 1.71 | 2.37 | 0.4508 | 3 | 6 |
| YA vs. MA | 2 | 5 | -8.06 to 2.06 | 2.31 | 0.4779 | 6 | 2 |
| YA vs. OA | 2 | 4 | -5.58 to 1.58 | 2.18 | 0.5377 | 6 | 6 |
| MA vs. OA | 5 | 4 | -4.06 to 6.06 | 0.77 | 0.9825 | 2 | 6 |
| **250 microns from soma** | **Predicted (LS) mean 1** | **Predicted (LS) mean 2** | **95.00% CI of diff.** | **q** | **p-value** | **N1** | **N2** |
| PW vs. PB | 2.29 | 1 | -3.68 to 6.25 | 1.01 | 0.9533 | 7 | 2 |
| PW vs. YA | 2.29 | 1.33 | -3.32 to 5.23 | 0.868 | 0.9728 | 7 | 3 |
| PW vs. MA | 2.29 | 2 | -6.34 to 6.91 | 0.168 | >0.9999 | 7 | 1 |
| PW vs. OA | 2.29 | 2 | -3.60 to 4.17 | 0.287 | 0.9996 | 7 | 4 |
| PB vs. YA | 1 | 1.33 | -5.99 to 5.32 | 0.23 | 0.9998 | 2 | 3 |
| PB vs. MA | 1 | 2 | -8.59 to 6.59 | 0.513 | 0.9963 | 2 | 1 |
| PB vs. OA | 1 | 2 | -6.36 to 4.36 | 0.726 | 0.986 | 2 | 4 |
| YA vs. MA | 1.33 | 2 | -7.82 to 6.49 | 0.363 | 0.999 | 3 | 1 |
| YA vs. OA | 1.33 | 2 | -5.40 to 4.06 | 0.549 | 0.9952 | 3 | 4 |
| MA vs. OA | 2 | 2 | -6.93 to 6.93 | 0 | >0.9999 | 1 | 4 |
|  | ^1^LS = Least Squares means used for unbalanced data | | | | | | |
|  | ^2^q = studentized range statistic | | | | | | |
|  | ^3^ p-value adjusted with a Tukey correction | | | | | | |
